# Heat Stress-induced Activation of MAPK Pathway Attenuates Atf1-dependent Epigenetic Inheritance of Heterochromatin in Fission Yeast

**DOI:** 10.1101/2023.07.06.547995

**Authors:** Li Sun, Yamei Wang, Quan-wen Jin

## Abstract

Eukaryotic cells are constantly exposed to various environmental stimuli. It remains largely unexplored how environmental cues bring about epigenetic fluctuations and affect heterochromatin stability. In the fission yeast *Schizosaccharomyces pombe*, heterochromatic silencing is quite stable at pericentromeres but unstable at the mating-type (*mat*) locus under chronic heat stress, although both loci are within the major constitutive heterochromatin regions. Here, we found that the compromised gene silencing at the *mat* locus at elevated temperature is linked to the phosphorylation status of Atf1, a member of the ATF/CREB superfamily. Constitutive activation of MAPK signaling disrupts epigenetic maintenance of heterochromatin at the *mat* locus even under normal temperature. Mechanistically, phosphorylation of Atf1 impairs its interaction with heterochromatin protein Swi6^HP1^, resulting in lower site-specific Swi6^HP1^ enrichment. Expression of non-phosphorylatable Atf1, tethering Swi6^HP1^ to the *mat3M*-flanking site or absence of the anti-silencing factor Epe1 can largely or partially rescue heat stress-induced defective heterochromatic maintenance at the *mat* locus.

## INTRODUCTION

Eukaryotic genomes contain two types of chromatin, namely euchromatin and heterochromatin, which are characterized according to their structure and compaction state, and the latter is crucial for regulating the gene expression pattern, cell differentiation and maintaining genomic stability (Allshire and Madhani, 2018; Bloom, 2014). In the fission yeast *Schizosaccharomyces pombe*, RNAi machinery contributes to the establishment of major constitutive heterochromatin at pericentromeres, telomeres and the silent mating-type region (*mat* locus) (Martienssen and Moazed, 2015; Volpe et al., 2002). In general, transcripts from heterochromatic regions, such as pericentromeric repeats, were processed into double strand small interfering RNAs (siRNAs) by RNase Dicer (Dcr1 in fission yeast) (Colmenares et al., 2007), then siRNAs were loaded to Argonaute (Ago1) to finally form functional RNAi-induced transcriptional silencing (RITS) complex, only containing single-stranded siRNAs (Verdel et al., 2004). The RITS complex can target nascent noncoding RNAs from heterochromatic regions through the single-stranded guide siRNAs and subsequently recruit the H3K9 methyltransferase Clr4 to establish H3K9me2/3 (Bayne et al., 2010; Hong et al., 2005), which can be bound by heterochromatin protein Swi6 and Chp2 through the conserved N-terminal chromo-domain (CD) (Jacobs and Khorasanizadeh, 2002; Jacobs et al., 2001; Maison and Almouzni, 2004). Heterochromatin proteins act as a platform to recruit downstream heterochromatin factors, such as the histone deacetylase (HDAC) Clr3 (Motamedi et al., 2008; Sugiyama et al., 2007), to initiate heterochromatin assembly. Once established, H3K9me2/3 can be firmly inherited independent of the mechanisms of heterochromatin establishment (Allshire and Madhani, 2018).

Heterochromatin plays an essential role in epigenetic gene silencing in organisms ranging from yeast to humans. Epigenetic states of heterochromatin can be stably inherited, but they are also reversible, which is true for not only facultative but also constitutive heterochromatin, and it can be influenced by environmental cues and thus evokes phenotypic variations. Eukaryotic cells are constantly exposed to various environmental stimuli, such as changes in osmotic pressure, oxygen, and temperature. Although the possible impact of the environment on epigenetic regulation has attracted considerable interest, it remains largely unknown how environmental cues bring about epigenetic fluctuations. So far, sporadic studies have demonstrated that heat stress is one of the most prevalent environmental stresses that trigger epigenetic alterations, which may negatively affect early embryonic development in mammals (Sun et al., 2023) and eye colour-controlling gene inactivation during early larval development in *Drosophila* (Seong et al., 2011), or serve as thermosensory input to positively control the rate of vernalization of the flowering plants after winter (Antoniou-Kourounioti et al., 2018; Feil and Fraga, 2012; Song et al., 2013).

In fission yeast, two redundant pathways contribute to establishment and maintenance of heterochromatin at the endogenous silent mating-type region. These two mechanisms rely on two major *cis* elements *cenH* and *REIII* acting as nucleation centers to recruit the H3K9 methyltransferase Clr4 via the RNAi machinery and the RNAi-independent ATF/CREB family proteins Atf1/ Pcr1, respectively (Hall et al., 2002; Jia et al., 2004a; Kim et al., 2004; Thon et al., 1999; Yamada et al., 2005). The initial nucleation and subsequent spreading of heterochromatin are further facilitated by Swi6^HP1^ and HDACs (including Clr3 and Clr6) (Jia et al., 2004a; Kim et al., 2004; Yamada et al., 2005). As two major stress-responsive transcription factors, it has been shown that Atf1 and Pcr1 are activated and regulated by Sty1, one of the mitogen-activated protein kinases (MAPKs), in response to high temperature, osmotic, oxidative and a number of other environmental stresses (Eshaghi et al., 2010; Lawrence et al., 2007; Reiter et al., 2008). Thus, it is plausible to assume that Atf1 and Pcr1 have the potential to render the heterochromatin stability at the silent mating-type region to be more resistant to ambient perturbations. However, contrary to this pre-assumption, recent studies demonstrated that the constitutive heterochromatin at centromeres is propagated stably whereas the epigenetic stability at the *mat* locus in vegetatively growing cells is sensitive to being continuously cultured at elevated temperatures (Greenstein et al., 2018; Nickels et al., 2022; Oberti et al., 2015). It has been established that the protein disaggregase Hsp104 is involved in buffering environmentally induced epigenetic variation at centromeres by dissolving cytoplasmic Dcr1 aggregates (Oberti et al., 2015). However, the reason for the absence of the buffering effect on heterochromatin at mating-type region under similar environmental stress remains elusive.

In this study, to explore the possible mechanism underlying the lack of epigenetic stability at the mating-type region in fission yeast under high temperature, we systematically performed genetic analyses combined with biochemical characterization. We found that heat stress-induced phosphorylation of Atf1 negatively influences its recruiting capability toward heterochromatin protein Swi6^HP1^, and thus it results in defective Atf1-dependent epigenetic maintenance of heterochromatin at the *mat* locus.

## RESULTS

### Gene silencing within constitutive heterochromatin at the *mat* locus is unstable under heat stress

To examine the stability of heterochromatin under environmental stresses, we first tested the robustness of heterochromatic silencing when cells were grown at 37℃, which is above the permissive temperature for *S. pombe* and causes acute temperature stress. We employed *S. pombe* strains with an *ura4^+^*reporter gene placed within two major constitutive heterochromatic regions, represented by pericentromere of chromosome I (*otr1R::ura4^+^*) and the mating-type region of chromosome II (*mat3M::ura4^+^*) (Ekwall et al., 1999; Thon and Klar, 1992) (Figure S1A). The silencing of the *ura4^+^*gene was monitored by poor colony formation on medium lacking uracil and vigorous growth on medium containing the counter-selective drug 5-fluoroorotic acid (5-FOA) (Figure S1B). Upon being grown at 37℃, we noticed obvious de-repression when the *ura4^+^* was inserted at *IR-R* element-proximal site within the mating-type region (*mat3M::ura4^+^*), but not at pericentromere (Figure S1B, C), indicating a mild loss of heterochromatic gene silencing at the *mat* locus under heat stress.

To confirm our above observations at high temperature, we used yeast strains in which an *ade6^+^* reporter gene was inserted into either the mating-type region (*mat3M::ade6^+^*) or the pericentromere (*otr1R::ade6^+^*) (Figure 1A). Cells with repressed *ade6^+^*within heterochromatic region gave rise to red colonies under limiting adenine conditions and failed to grow on medium without adenine, whereas de-repressed *ade6^+^* allowed cells to form white colonies or vigorous growth (Figure 1B). Consistent with previous study (Oberti et al., 2015), *otr1R::ade6^+^* cells formed almost fully red colonies at all tested temperatures (Figure 1B, C). In contrast, *mat3M::ade6^+^* cells gave rise to variegated colonies or white colonies with low degrees of redness, or even were able to grow on medium without adenine at 37℃ (Figure 1B, C). Accordingly, the mRNA levels of *ade6^+^*, as measured by quantitative RT-PCR, increased by 7-fold in *mat3M::ade6^+^* cells but only 2-fold in *otr1R::ade6^+^* cells at 37℃ compared to the same cells grown at 30℃ (Figure 1D). These results are consistent with recent reports that heterochromatin at pericentromere is largely maintained and that at the *mat* locus seems to be unstable under heat stress (Greenstein et al., 2018; Nickels et al., 2022; Oberti et al., 2015).

**Figure 1.**
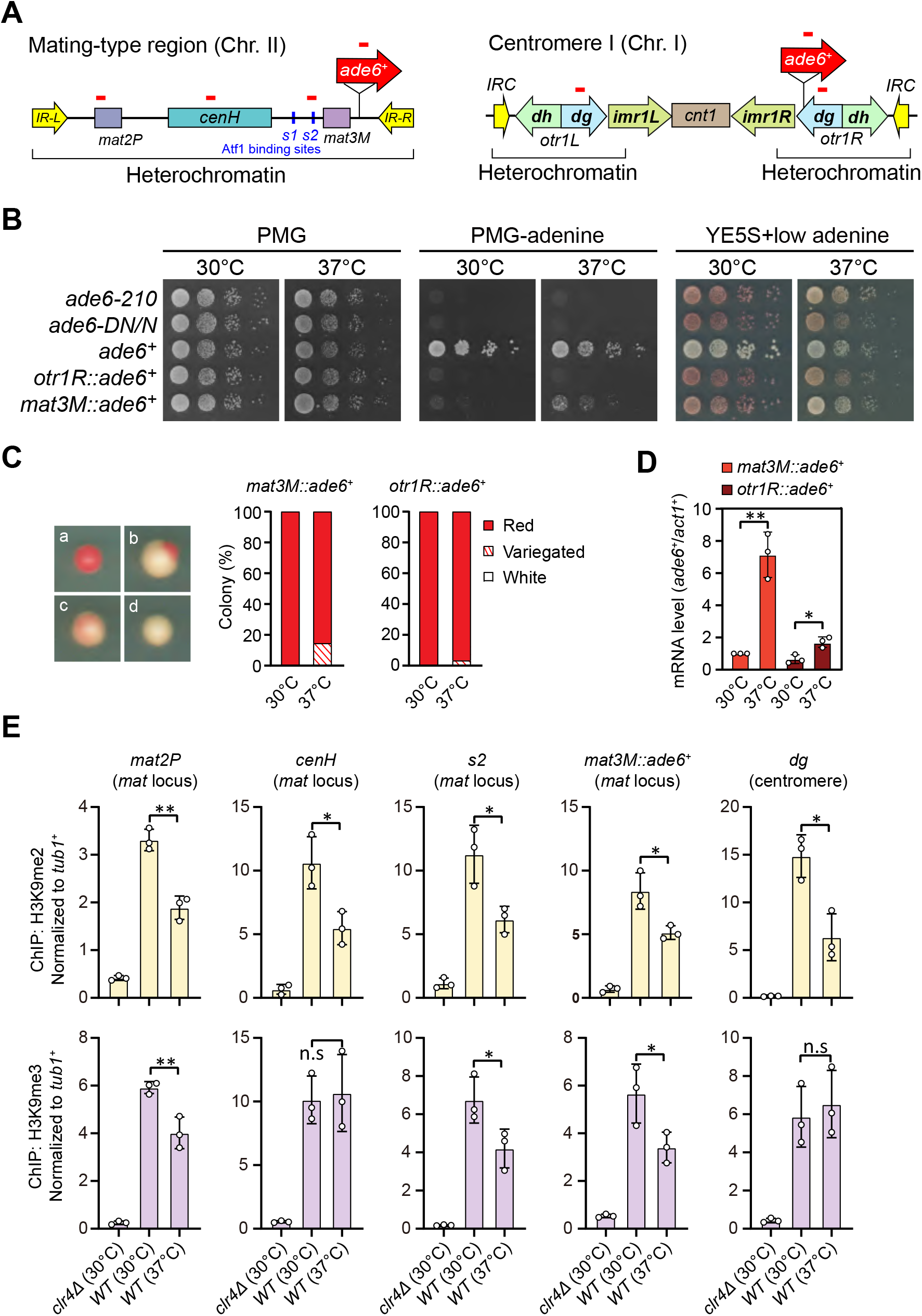
Heat stress leads to gene silencing defects at the mating-type region. (A) Schematic of an *ade6^+^* reporter gene inserted into mating-type region and pericentromeric region. Primer positions for RT-qPCR or ChIP analysis are indicated (red bars). *cenH*, a DNA element homologous to pericentromeric repeats; *mat2-P* and *mat3-M*, two silent cassettes used for mating-type switching; *IR-L* and *IR-R*, inverted repeats and boundary elements; *s1* and *s2*, two Atf1 binding sites; *cnt1*, central core; *imr1*, innermost repeats; *otr1*, outermost repeats; *dg* and *dh*, tandem repeats in *otr*; *IRC*, inverted repeats and boundary elements. (B) Expression of the *ade6^+^* reporter monitored by serial dilution spot assay at 30°C and 37°C. The media used were nonselective PMG, selective PMG without adenine and YE5S with low concentration of adenine. (C) Expression of the *ade6^+^* reporter monitored by colony color assay. (Left) Representative colonies of *ade6^+^* reporter fully repressed (red), partially repressed (variegated) and completely expressed (white) on low adenine medium; (Right) Variegated colonies were quantified at 30°C and 37°C. *n* > 500 colonies counted for each sample. (D) RT-qPCR analyses of *ade6^+^* reporter. The relative *ade6^+^* mRNA level was quantified with a ratio between *mat3M::ade6^+^* and *act1^+^* in 30°C samples being set as 1.00. Error bars indicate mean ± standard deviation of three independent experiments. Two-tailed unpaired *t*-test was used to derive *p* values. n.s, not significant; \**p*<0.05; \*\**p*<0.01; \*\*\**p*<0.001. (E) ChIP-qPCR analyses of H3K9me2/3 levels at heterochromatic loci. Relative enrichment of H3K9me2/3 was normalized to that of a *tub1^+^* fragment. Error bars represent standard deviation of three experiments. Two-tailed unpaired *t*-test was used to derive *p* values. n.s, not significant; \**p*<0.05; \*\**p*<0.01; \*\*\**p*<0.001.

We also examined the mRNA and translated protein levels of a *gfp^+^* transgene inserted at *mat3M* locus (*mat3M::gfp^+^*) or pericentromeric repeat region (*imr1R::gfp^+^*) (Figure S2A). Consistent with our results in *mat3M::ade6^+^* cells, both *gfp^+^* mRNA levels and GFP protein levels were increased significantly in *mat3M::gfp^+^* cells, but not in *imr1R::gfp^+^*cells at 37℃ (Figure S2B). We noticed that the GFP levels were actually slightly decreased in *imr1R::gfp^+^* cells at 37℃, which might be due to the instability of the GFP under heat stress (Figure S2B).

To further examine whether our observed heterochromatic gene silencing defects at *mat3M* under heat stress is coupled to compromised maintenance of heterochromatin, we performed ChIP followed by quantitative PCR (ChIP-qPCR) to monitor the levels of H3K9me2 and H3K9me3, the hallmarks of heterochromatin. Our results showed that when cells were grown at 37℃, H3K9me2 was reduced modestly at all heterochromatic regions, and intriguingly, H3K9me3 enrichment was similarly reduced within *cenH* element-surrounding regions except pericentromeric repeats and the *cenH* region itself (Figure 1E), which shares homology to pericentromeric repeats and is required for nucleation of heterochromatin at the *mat* locus. This is consistent with recent finding that H3K9me3, but not H3K9me2, is a more reliable hallmark for heterochromatin (Cutter DiPiazza et al., 2021; Jih et al., 2017). Together, these results demonstrated that gene silencing and heterochromatin at the *mat* locus is much more sensitive to high temperature than pericentromeric regions in fission yeast.

### Heat stress compromises reestablishment of a stable epigenetic state of heterochromatin at the *mat* locus

One previous study has revealed that the RNAi mechanism is still functional and actively confers the robustness of epigenetic maintenance of heterochromatin at centromere upon heat stress (Oberti et al., 2015). At the *mat* locus, RNAi mechanism is similarly required for the nucleation of heterochromatin at the *cenH* element and subsequent spreading across the entire *mat* locus, but it is dispensable for the maintenance of heterochromatin (Jia et al., 2004a; Kim et al., 2004). Our observation that H3K9me3 was largely maintained within *cenH* at 37℃ (Figure 1E) prompted us to investigate whether the defective maintenance of heterochromatin at *mat3M* might be masked by RNAi-mediated *de novo* heterochromatin assembly. Indeed, in accordance with our assumption, the variegated colonies from *mat3M::ade6^+^* cells at 37℃ restored gene silencing rapidly at normal temperature 30℃ after being re-plated on medium containing limited adenine (Figure 2A, B). However, when this re-plating assay was applied to *dcr1Δ*, one of the RNAi mutants, a considerable proportion of cells still emerged as variegated colonies (designated as *dcr1Δ^V^*), which was in sharp contrast to wild type cells (Figure 2B). Our RT-qPCR analyses confirmed that the mRNA levels of the reporter *ade6^+^* increased dramatically in *dcr1Δ^V^* cells compared to those in *dcr1Δ^R^* (refers to “red” colonies) cells at 30℃. Furthermore, the de-repression also correlated with severe reduction in H3K9me3 levels within the entire *mat* locus in *dcr1Δ^V^* cells (Figure 2C, D). These data strongly suggested that heat stress also leads to defective reestablishment of stable heterochromatin at the *mat* locus.

**Figure 2.**
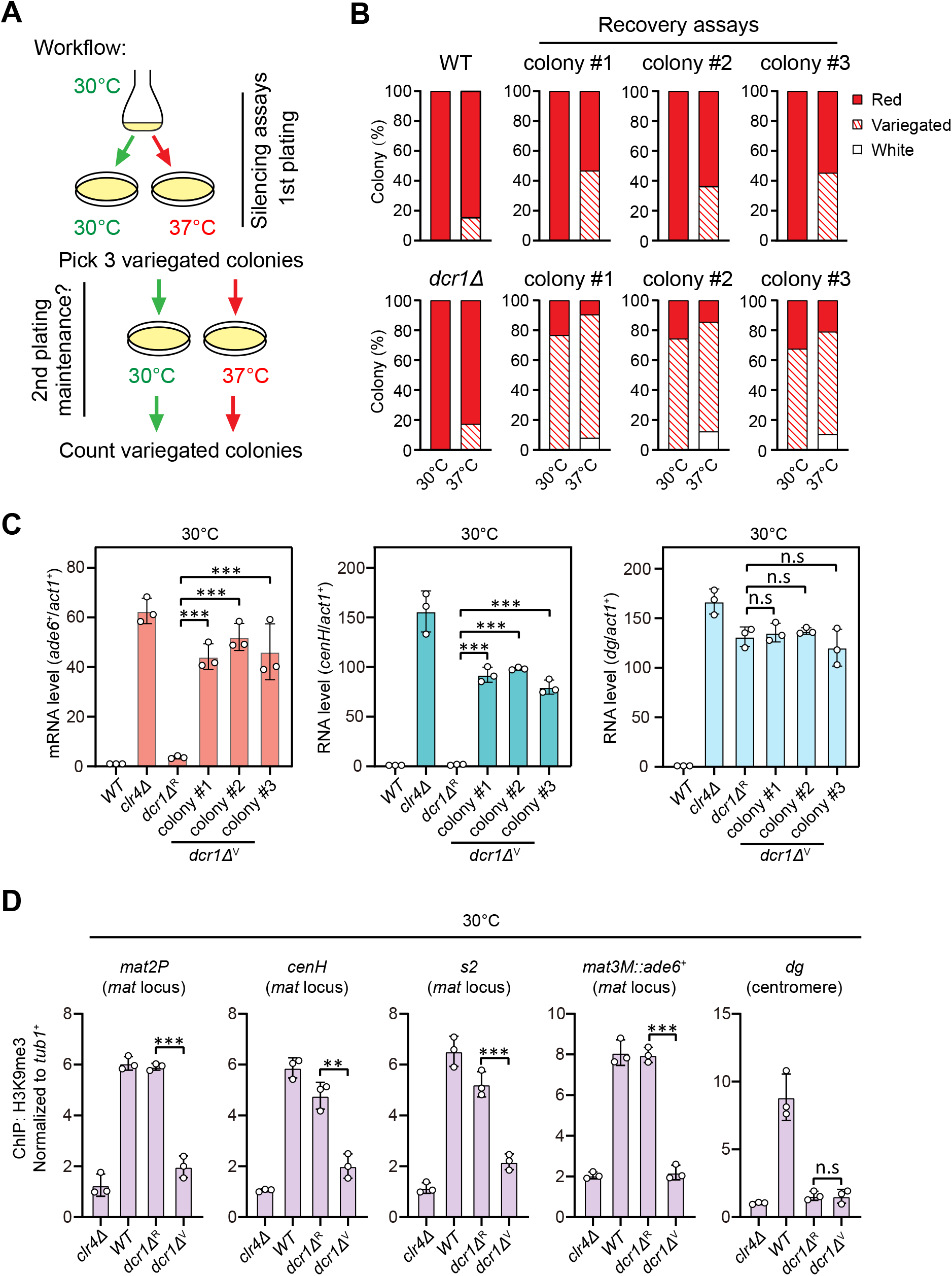
Heat stress compromises reestablishment of a stable epigenetic state of heterochromatin at the mating-type region. (A) Workflow of gene silencing recovery assays. 1st plating: strains were plated on low adenine medium at 30°C and 37°C. 2nd plating: three variegated colonies (*ade6^+^* was partially repressed) from low adenine plates at 37°C were collected, resuspended in water and then directly re-plated on low adenine medium and grown at 30°C or 37°C. Variegated colonies were counted to assess the gene silencing defect. (B) Quantified results of *ade6^+^*gene silencing recovery assays. Variegated colonies on low adenine medium from 1st plating and 2nd plating were counted. *n* > 500 colonies counted for each sample. (C) RT-qPCR analyses of *ade6^+^* reporter, *cenH* and *dg* transcripts. *dcr1Δ*^R^ and *dcr1Δ*^V^ indicate red colonies and variegated colonies respectively when *dcr1Δ* cells were grown on low adenine plate at 37°C. Three variegated colonies were picked and re-plated on low adenine medium and grown at 30°C. The relative transcript level was quantified with a ratio between respective transcript and *act1^+^* in 30°C wild type samples being set as 1.00. Error bars indicate mean ± standard deviation of three independent experiments. Two-tailed unpaired *t*-test was used to derive *p* values. n.s, not significant; \**p*<0.05; \*\**p*<0.01; \*\*\**p*<0.001. (D) ChIP-qPCR analyses of H3K9me3 levels at heterochromatic loci. Samples were collected as in (C). Relative enrichment of H3K9me3 was normalized to that of a *tub1^+^* fragment. Error bars represent standard deviation of three experiments. Two-tailed unpaired *t*-test was used to derive *p* values. n.s, not significant; \**p*<0.05; \*\**p*<0.01; \*\*\**p*<0.001.

### Phosphorylation of Atf1 causes heat stress-induced defective epigenetic maintenance of the *mat* locus

It has been recently established that a composite DNA element within *REIII* at the *mat* locus contains binding sequences for Atf1/Pcr1, Deb1 and the origin recognition complex (ORC), which act together in epigenetic maintenance of heterochromatin in the absence of RNAi nucleation with Atf1 as the dominating contributor (Wang et al., 2021). Two previous studies showed that extracellular stresses induce phosphorylation of *Drosophila* dATF-2 and mouse ATF7, two homologs of *S. pombe* Atf1, this provokes their release from heterochromatin and thus disrupted heterochromatic maintenance (Liu et al., 2019; Seong et al., 2011). Given that Atf1 can be phosphorylated by MAPK under heat stress (Samejima et al., 1997; Shiozaki et al., 1998), we surmised that heat stress-induced phosphorylation of Atf1 might similarly cause its release from the *mat* locus in fission yeast. Surprisingly, ChIP-qPCR analyses showed that Atf1 abundance at the *mat* locus was not altered at 37℃, although it was indeed increased modestly at *SPCC320.03*, an euchromatic target of Atf1/Pcr1 (Eshaghi et al., 2010) (Figure 3A).

**Figure 3.**
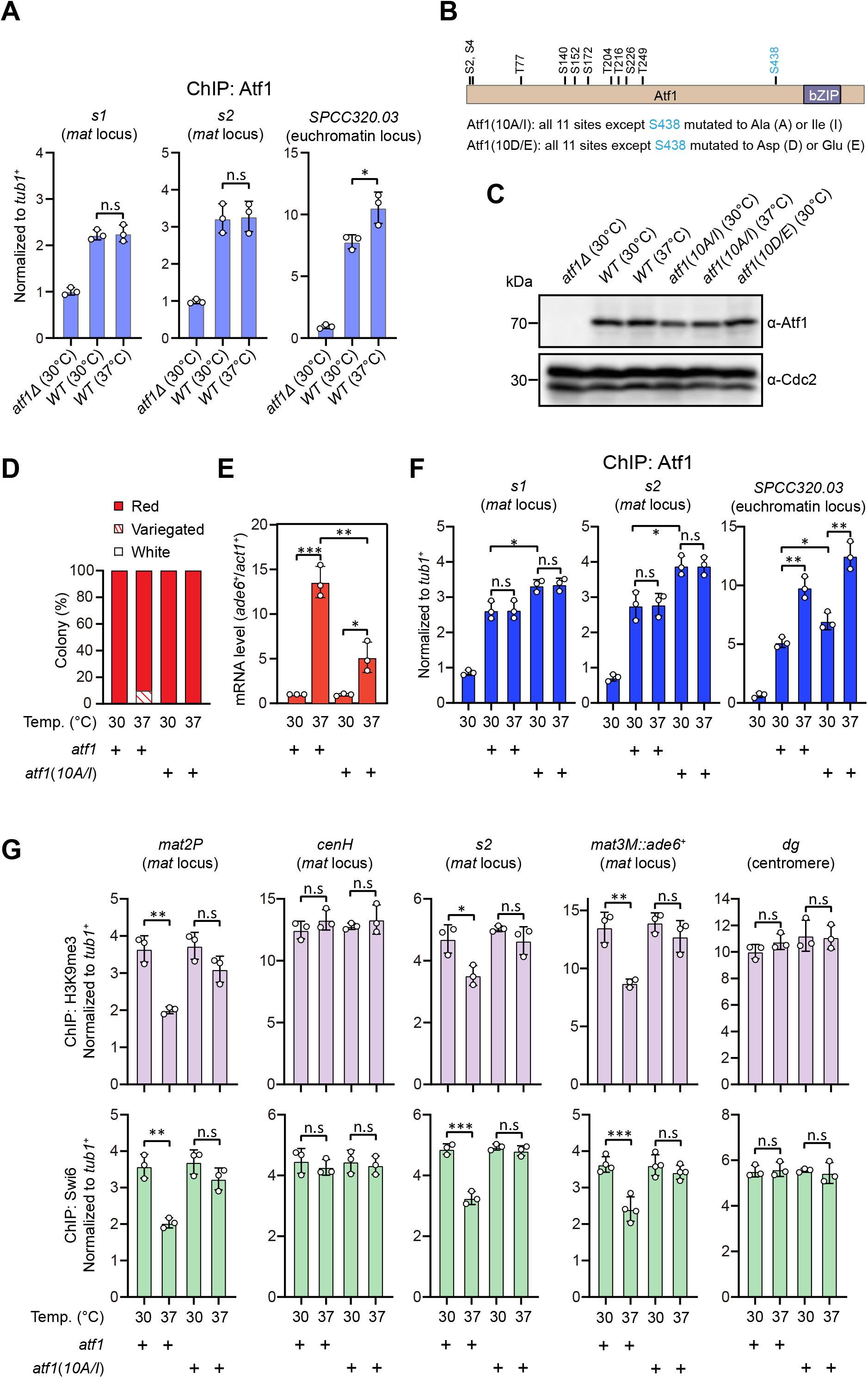
Heat stress-induced defective heterochromatic maintenance at the mating-type region can be rescued by non-phosphorylatable Atf1(10A/I). (A) ChIP-qPCR analyses of Atf1 levels at two Atf1 binding sites (*s1*, *s2*) within mating-type region and an euchromatic target of Atf1 (SPCC320.03). Relative enrichment of Atf1 is normalized to that of a *tub1^+^* fragment. Error bars represent standard deviation of three experiments. Two-tailed unpaired *t*-test was used to derive *p* values. n.s, not significant; \**p*<0.05; \*\**p*<0.01; \*\*\**p*<0.001. (B) Schematic depiction of the Atf1 protein with the substitutions of the 10 putative phosphorylation sites to alanines or isoleucines (10A/I), or aspartic acids or glutamic acids (10D/E) indicated. (C) Western blotting analyses of the protein level of Atf1 in *atf1Δ* background cells expressing HA-Atf1, HA-Atf1(10A/I) or HA-Atf1(10D/E) under the control of *sty1^+^* promoter. (D) Expression of the *mat3M::ade6^+^* reporter monitored by colony color assay in *atf1Δ* cells expressing *P_sty1_-HA-atf1* or *P_sty1_-HA-atf1(10A/I)* as in Figure 1C. *n* > 500 colonies counted for each sample. (E) RT-qPCR analyses of the *mat3M::ade6^+^* reporter in *atf1Δ* cells expressing *P_sty1_-HA-atf1* or *P_sty1_-HA-atf1(10A/I)*. (F) ChIP-qPCR analyses of Atf1 levels at two Atf1 binding sites within mating-type region and an euchromatic target of Atf1 (SPCC320.03) in *atf1Δ* cells expressing *P_sty1_-HA-atf1* or *P_sty1_-HA-atf1(10A/I)*. (G) ChIP-qPCR analyses of H3K9me3 and Swi6 levels at heterochromatic loci in *atf1Δ* cells expressing *P_sty1_-HA-atf1* or *P_sty1_-HA-atf1(10A/I)*.

Atf1 contains 11 putative MAPK phosphorylation sites, 10 out of them are within the first half of Atf1 (Lawrence et al., 2007) (Figure 3B). Thus, we asked whether the phosphorylation status of Atf1 is causative to defective maintenance of heterochromatin at the *mat* locus under heat stress. To test this, we constructed strains carrying ectopically expressed *HA-atf1*, *HA-atf1(10A/I)* or *HA-atf1(10D/E)* under the control of the *sty1^+^* promoter (*P_sty1_*) with the endogenous *atf1^+^* gene deleted (Salat-Canela et al., 2017). Alleles of *atf1(10A/I)* and *HA-atf1(10D/E)* harbor 10 non-phosphorylatable alanines (Ala, A) and isoleucines (Ile, I), or phosphomimetic aspartic acids (Asp, D) and glutamic acids (Glu, E) replacing serines or threonines, respectively (Salat-Canela et al., 2017). Our immunoblotting analyses showed that the protein levels of HA-Atf1 and HA-Atf1(10A/I) were comparable (Figure 3C). Intriguingly, cells expressing *P_sty1_*-*HA-atf1(10A/I)* fully rescued the gene silencing defects observed in *P_sty1_*-*HA-atf1* cells at 37℃, which was confirmed by RT-qPCR analyses (Figure 3D, E). Consistently, more Atf1 was maintained at the *mat* locus in *P_sty1_*-*HA-atf1(10A/I)* cells compared to that in *P_sty1_*-*HA-atf1* cells grown at both 30℃ and 37℃ (Figure 3F). Moreover, ChIP-qPCR analyses showed that the enrichment of H3K9me3 and heterochromatin protein Swi6^HP1^ at the *mat* locus in *P_sty1_*-*HA-atf1(10A/I)* cells was also restored to the level of wild type cells (Figure 3G). To our surprise, cells expressing *P_sty1_*-*HA-atf1(10D/E)* were almost completely unable to grow at 37℃ (Figure S3), which may be due to the toxicity caused by constitutive level of Atf1(10D/E). This hindered our further investigation on the effect of Atf1(10D/E) on maintenance of heterochromatin at the *mat* locus under heat stress. Taken together, these data demonstrated that phosphorylation of Atf1 causes heat stress-induced defective epigenetic maintenance of the *mat* locus.

### Heat stress-induced phosphorylation of Atf1 reduces its binding affinity to Swi6^HP1^

Previous studies have shown that Atf1 contributes to the epigenetic maintenance of the *mat* locus by actively recruiting the H3K9 methyltransferase Clr4, heterochromatin protein Swi6, and two HDACs Clr3 and Clr6 (Jia et al., 2004a; Kim et al., 2004). To explore whether phosphorylation of Atf1 compromises its capability of recruiting these heterochromatic factors at the *mat* locus under heat stress, we performed *in vitro* pull-down assays using yeast lysates prepared from cultures grown at either 30℃ or 37℃ and bacterially expressed GST-Clr3, GST-Clr4, MBP-Clr6 and His-Swi6. We found that Atf1 from cells grown at 37℃ almost completely lost its binding to Swi6^HP1^ but not the other three heterochromatic proteins (Figure 4A and Figure S4A). Very interestingly, non-phosphorylatable Atf1(10A/I) from 37℃ cultures bound Swi6^HP1^ more efficiently than wild type Atf1, and phosphomimetic Atf1(10D/E) rendered weak binding to Swi6^HP1^ even when it was derived from 30℃ cultures (Figure 4A).

**Figure 4.**
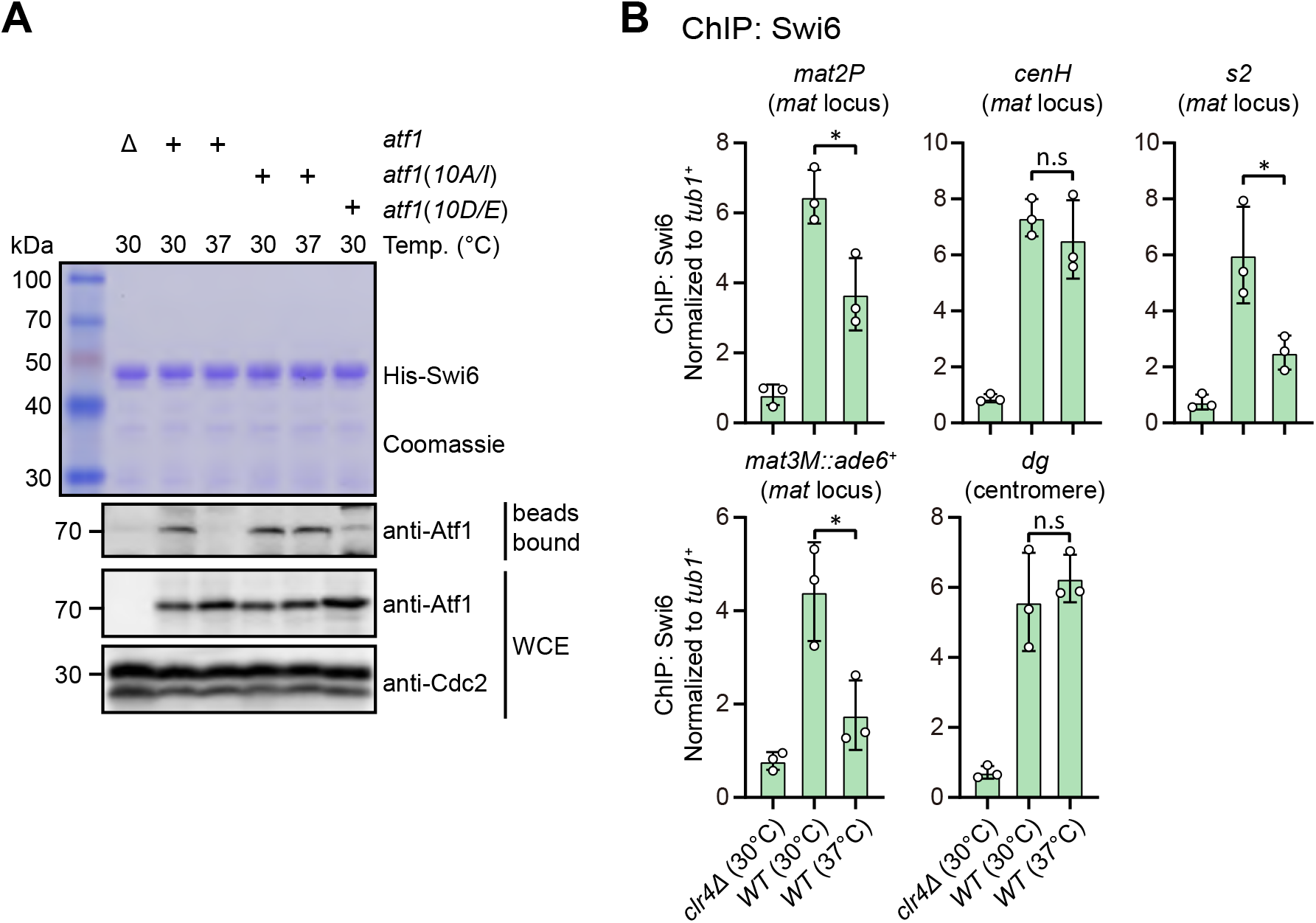
Phosphorylation of Atf1 impairs its interaction with Swi6^HP1^. (A) Binding affinity between Atf1 and Swi6^HP1^ is maintained for non-phosphorylatable Atf1(10A/I) at 37°C, but disrupted for phosphomimetic Atf1(10D/E) even at 30°C. Yeast lysates from *atf1Δ* cells expressing *P_sty1_-HA-atf1*, *P_sty1_-HA-atf1(10A/I)* or *P_sty1_-HA-atf1(10D/E)* grown at either 30°C or 37°C were incubated with bacteria-expressed 6His-Swi6 in *in vitro* pull-down assays. Bound and total Atf1 were detected by immunoblotting with Cdc2 used as a loading control. Results are representative of three independent experiments. (B) ChIP-qPCR analyses of Swi6 levels at heterochromatic loci. Relative enrichment of Swi6 was normalized to that of a *tub1^+^* fragment. Error bars represent standard deviation of three experiments. Two-tailed unpaired *t*-test was used to derive *p* values. n.s, not significant; \**p*<0.05; \*\**p*<0.01; \*\*\**p*<0.001.

Our ChIP-qPCR analyses confirmed that Swi6^HP1^ enrichment was indeed reduced at the *mat* locus, but not at pericentromeric repeats under heat stress (Figure 4B). In addition, we also noticed that Clr3 level was slightly decreased at regions distal to the *cenH* nucleation center (i.e. *mat2P* and *mat3M*) but not at pericentromeres and the *cenH* under heat stress (Figure S4B). This was in sharp contrast to the actually slight increase of the binding between Clr3 and Atf1 at 37℃ detected by *in vitro* pull-down assays (Figure S4A). Consistent with our *in vitro* pull-down assays, Clr4 and Clr6 enrichment was not altered at all tested heterochromatin sites under heat stress (Figure S4C, D).

Together, all these results suggested that most likely phosphorylation of Atf1 induced by heat stress disrupts its binding affinity to Swi6^HP1^ and thus attenuates Swi6^HP1^ abundance at the *mat* locus, this consequently causes the defective maintenance of heterochromatin specifically at this site.

### Constitutive activation of MAPK signaling leads to defective epigenetic maintenance of heterochromatin at the *mat* locus under normal temperature

It has been well established that the MAPK Sty1 is constitutively activated in *wis1-DD* mutant (carrying S469D and T473D mutations), and Sty1 phosphorylates and activates Atf1 in response to high temperature (Eshaghi et al., 2010; Lawrence et al., 2007; Reiter et al., 2008). It is fairly possible that permanent activation of Sty1 may also lead to defective epigenetic maintenance of heterochromatin at the *mat* locus even at 30℃. To test this possibility, we employed a yeast strain in which *cenH* site was replaced with an *ade6^+^* reporter gene (*kΔ::ade6^+^*) (Grewal and Klar, 1996; Thon and Friis, 1997) to remove RNAi-mediated heterochromatin nucleation (Figure 5A), and *kΔ::ade6^+^*displays one of two distinct statuses: being expressed (*ade6*-on) or being silenced (*ade6*-off). In *ade6*-off cells, Atf1 becomes the major determinant factor for epigenetic maintenance of heterochromatin at the *mat* locus (Wang and Moazed, 2017; Wang et al., 2021). Notably, cells expressing two copies, but not one copy, of *wis1-DD* showed severe gene silencing defects (Figure 5B), and consistently the mRNA levels of the *kΔ::ade6^+^* increased dramatically in these cells (Figure 5C). Our immunoblotting results confirmed that the levels of both the active form of Sty1 (i.e. phosphorylated of Sty1) and its downstream effector Atf1 increased in *wis1-DD* mutants regardless of its copy number (Figure 5D), indicating constitutive activation of the Wis1/Sty1-mediated MAPK signaling. Furthermore, *in vitro* pull-down assays demonstrated that the interaction between Atf1 and Swi6^HP1^ was also largely disrupted in *wis1-DD* mutants (Figure 5E). Consistently, ChIP-qPCR analyses showed that the abundance of both H3K9me3 and Swi6^HP1^ bound at the *mat* locus but not at pericentromere decreased dramatically in cells with two copies of *wis1-DD* (Figure 5F). Therefore, our data lent support to the idea that constitutive activation of MAPK signaling and resulted Atf1 phosphorylation can also eventually lead to defective epigenetic maintenance and inheritance of heterochromatin at the *mat* locus under normal temperature.

**Figure 5.**
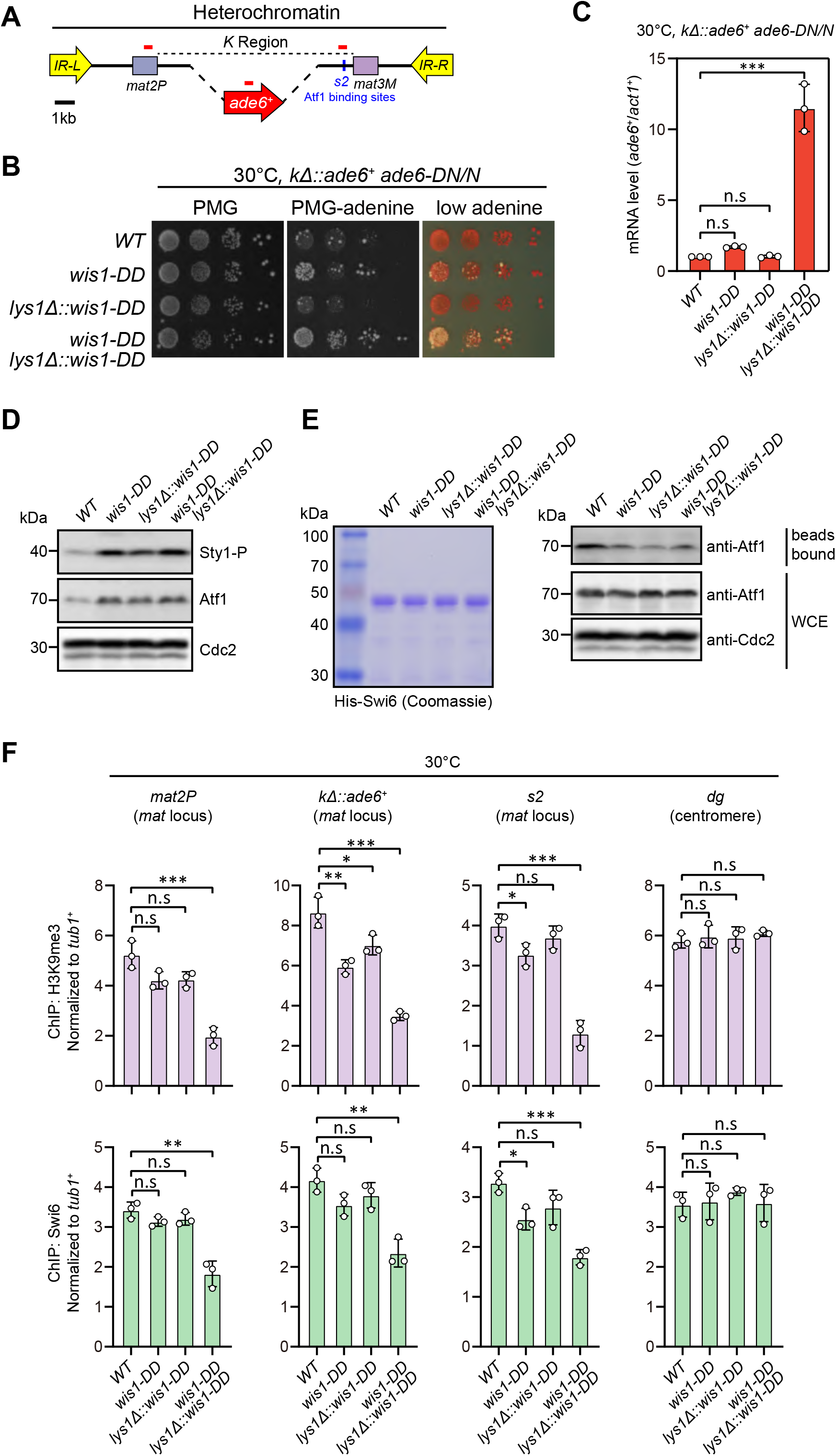
Constitutive activation of MAPK signaling pathway leads to defective epigenetic maintenance of heterochromatin at the mating-type region. (A) Schematic of the mating-type region in *kΔ::ade6*^+^ strain. A 7.5kb DNA sequence (*K* region) between *mat2P* and *mat3M* locus was replaced with *ade6^+^* reporter. Primer positions for RT-qPCR or ChIP analysis are indicated (red bars). (B) Expression of the *kΔ::ade6^+^* reporter monitored by serial dilution spot assay at 30°C as in Figure 1B. Constitutive activation of one of the MAPK signaling pathways was achieved by expressing *wis1-DD* (*wis1-S469D;T473D*) mutant at either endogenous locus or ectopically at *lys1+* (*lys1Δ::wis1-DD*) or simultaneously at both loci. (C) RT-qPCR analyses of the *kΔ::ade6^+^* reporter. (D) Western blotting analyses of the phosphorylated Sty1 and the total protein of Atf1. (E) Binding affinity between Atf1 and Swi6^HP1^ was detected by *in vitro* pull-down assays as in Figure 4A. Yeast lysates were prepared from wild type or *wis1-DD* cells grown at 30°C. Results are representative of three independent experiments. (F) ChIP-qPCR analyses of H3K9me3 and Swi6 levels at heterochromatic loci in wild type and *wis1-DD* cells grown at 30°C.

### Tethering Swi6^HP1^ to the *mat* locus rescues heat stress-induced defective heterochromatic maintenance at the *mat* locus

Our above results demonstrated that lowered Swi6^HP1^ abundance at the *mat* locus under heat stress is very likely the major cause for defective heterochromatin stability. To further test this assumption, we adopted an artificial tethering system involving bacterial tetracycline operator sequence (*tetO*) and repressor protein (TetR^off^) (Bayne et al., 2010; Ragunathan et al., 2015). A sequence containing four *tetO* upstream of *ade6^+^* reporter gene was inserted into *mat3M* locus (*mat3M::4xtetO-ade6^+^*), and Swi6 lacking its chromo domain (CD) was fused with TetR^off^ (TetR^off^-Swi6^ΔCD^), which should allow the fusion protein to bind specifically at *4xtetO-ade6^+^* (Figure 6A). This indeed resulted in efficient recruitment of TetR^off^-Swi6^ΔCD^ to the reporter at both 30℃ and 37℃ (Figure 6B). And cells expressing TetR^off^-Swi6^ΔCD^ but not TetR^off^ or Swi6^ΔCD^ only rendered completely red colonies at 37℃ (Figure 6C), indicating full rescue of silencing defects at the *mat* locus under heat stress. Our RT-qPCR and ChIP-qPCR analyses showed that the mRNA level of the *ade6^+^* was decreased dramatically and H3K9me3 level was restored, respectively, in cells expressing TetR^off^-Swi6^ΔCD^ at 37℃ (Figure 6D, E), demonstrating that tethering Swi6^HP1^ to the *mat* locus was sufficient to rescue heat stress-induced defective epigenetic maintenance of heterochromatin. This also supports the idea that low Swi6^HP1^ affinity and abundance at the *mat* locus brought about by MAPK-dependent phosphorylation of Atf1 is the major contribution factor for loss of heterochromatin at higher temperatures.

**Figure 6.**
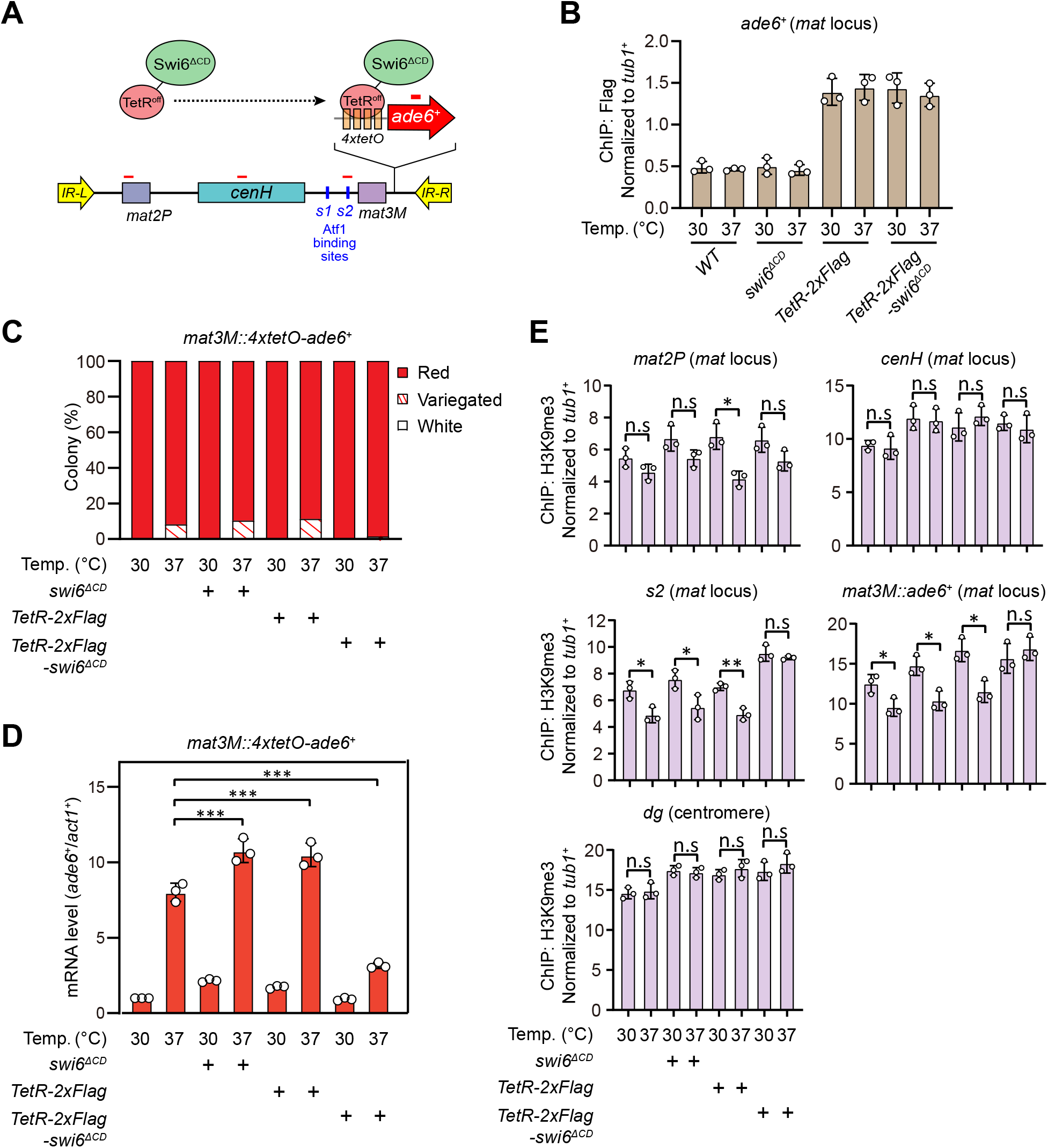
Tethering Swi6 ^HP1^ to the *mat3M*-flanking site rescues heat stress-induced defective epigenetic maintenance of heterochromatin at the *mat* locus. (A) Schematic of tethering Swi6^HP1^ to the *mat* locus. A sequence containing four tetracycline operators located upstream of *ade6^+^* reporter gene (*4xtetO-ade6^+^*) was inserted next to *mat3M* locus, and Swi6 lacking CD domain was fused with TetR^off^ (TetR^off^-Swi6^ΔCD^) and a 2xFlag tag. Primer positions for RT-qPCR or chromatin immunoprecipitation (ChIP) analysis are indicated (red bars). (B) ChIP-qPCR analyses of Flag-tagged TetR^off^-Swi6^ΔCD^ at *4xtetO-ade6^+^* locus. Relative enrichment of TetR^off^-Swi6^ΔCD^ was normalized to that of a *tub1^+^* fragment. Error bars represent standard deviation of three experiments. (C) Expression of the *mat3M::4xtetO-ade6^+^* reporter monitored by colony color assay. *n* > 500 colonies counted for each sample. (D) RT-qPCR analyses of the *mat3M::4xtetO-ade6^+^* reporter. (E) ChIP-qPCR analyses of H3K9me3 levels at heterochromatic loci in Swi6^HP1^-tethered cells.

### Increased heterochromatin spreading in *epe1Δ* alleviates silencing defects at the *mat* locus upon heat stress

At the *mat* locus, both HDACs Sir2 and Clr3 contribute to heterochromatin spreading in concert with RNAi-directed heterochromatin nucleation (Shankaranarayana et al., 2003; Yamada et al., 2005). We noticed that Clr3 level was slightly decreased at regions flanking the *cenH* nucleation center but not at the *cenH* itself at high temperature (Figure S4B). In *S. pombe*, several factors have been identified to be required for preventing uncontrolled heterochromatin spreading and massive ectopic heterochromatin, notably the JmjC domain-containing protein Epe1 (Braun et al., 2011; Wang et al., 2013; Zofall and Grewal, 2006), the histone acetyltransferase Mst2 (Wang et al., 2015) and the transcription elongation complex Paf1C, which includes five subunits Paf1, Leo1, Cdc73, Prf1 and Tpr1 (Kowalik et al., 2015; Sadeghi et al., 2015). If the heterochromatin defects at the *mat* locus under heat stress are also due to the compromised spreading mediated by Clr3, then we would expect that a genetic background that is more permissive to heterochromatin spreading might overcome this barrier and rescue silencing defects. Intriguingly, we found that *epe1Δ*, but not *mst2Δ* or *leo1Δ*, indeed moderately rescued the silencing defects at the *mat* locus based on our silencing assays and RT-qPCR analyses of the *mat3M::ade6^+^* transcripts (Figure 7A, B). Moreover, ChIP-qPCR analysis also showed that H3K9me3 level at the *mat* locus in *epe1Δ* cells was more robust than wild type, *mst2Δ* or *leo1Δ* cells at 37℃ (Figure 7C).

**Figure 7.**
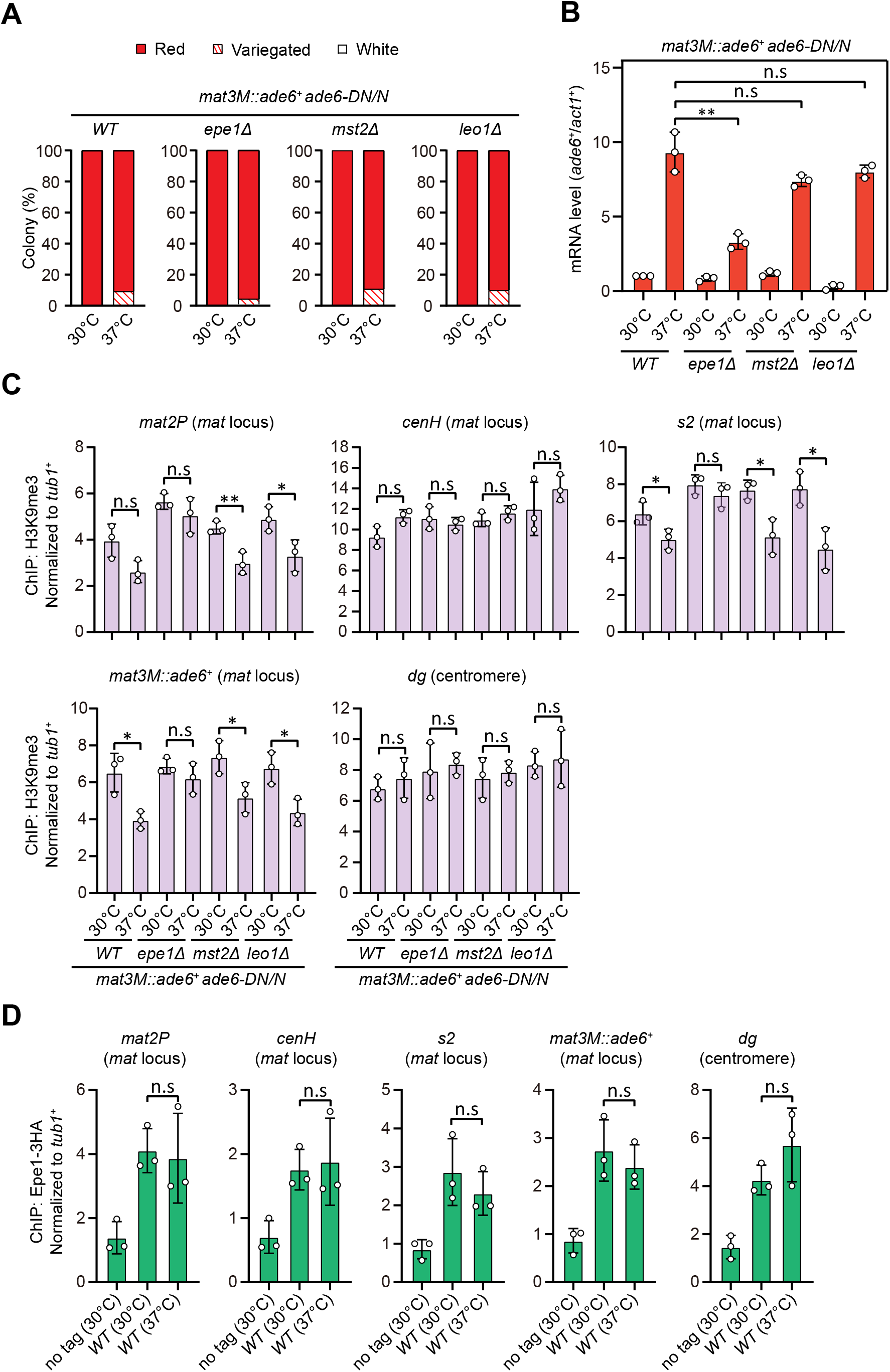
Deletion of anti-silencing factor Epe1 rescues heat stress-induced defective epigenetic maintenance of heterochromatin at mating-type region. (A) Expression of the *mat3M::ade6^+^* reporter monitored by colony color assay in *epe1Δ, mst2Δ* or *leo1Δ* cells. *n* > 500 colonies counted for each sample. (B) RT-qPCR analyses of the *mat3M::ade6^+^* reporter in *epe1Δ, mst2Δ* or *leo1Δ* cells. (C) ChIP-qPCR analyses of H3K9me3 levels at heterochromatic loci in *epe1Δ, mst2Δ* or *leo1Δ* cells. (D) ChIP-qPCR analyses of Epe1 levels at heterochromatic loci in wild type cells grown at 30°C or 37°C. Relative enrichment of Epe1-3HA was normalized to that of a *tub1^+^* fragment. Error bars represent standard deviation of three experiments. Two-tailed unpaired *t*-test was used to derive *p* values. n.s, not significant; \**p*<0.05; \*\**p*<0.01; \*\*\**p*<0.001.

Based on above data, we were suspicious that more Epe1 might enrich at the *mat* locus under heat stress to antagonize heterochromatic silencing. To test this possibility, we measured Epe1 abundance at the *mat* locus under heat stress by ChIP-qPCR. In contrast to our anticipation, the levels of Epe1 at the *mat* locus were similar in cells cultured at 30℃ and 37℃ (Figure 7D), indicating Epe1 is not actively involved in competition with Swi6^HP1^ at this site.

## DISCUSSION

Using fission yeast as the model organism, previous studies have shown that heterochromatic silencing at pericentromeres is largely stable under chronic heat stress conditions (Oberti et al., 2015), but heterochromatin becomes unstable and a significant derepression occurs at the mating-type region at elevated temperatures (Greenstein et al., 2018; Nickels et al., 2022), although both loci are within the major constitutive heterochromatin regions. It has been established that the temperature-insensitivity of centromeric heterochromatin is mainly attributed to the buffering effect of the protein disaggregase Hsp104, which actively prevents formation of Dicer aggregations in cytoplasmic inclusions and promotes the recycling of resolubilized Dcr1 (Oberti et al., 2015). However, the mechanistic details and the physiological relevance of the loss of heterochromatin stability at the *mat* locus under similar environmental stress are unknown.

### Instability of ATF/CREB family protein-dependent temperature-sensitive heterochromatin may operate through distinct mechanisms

In the current study, we revisited the unusual temperature-sensitive heterochromatin at the mating-type region in fission yeast. It is known that at this locus, both RNAi machinery and multiple factors, including transcription factors Atf1/Pcr1 and Deb1 and the origin recognition complex (ORC), are required for local heterochromatin formation (Greenstein et al., 2018; Jia et al., 2004a; Kim et al., 2004; Nickels et al., 2022; Thon et al., 1999; Wang et al., 2021; Yamada et al., 2005). Among these factors, Atf1/Pcr1 heterodimer binds to the *cis* elements flanking the *mat3M* cassette, which are mainly composed of closely juxtaposed 137bp DNA sequence containing *s1* and *REIII* elements (Jia et al., 2004a; Kim et al., 2004; Nickels et al., 2022; Thon et al., 1999; Wang et al., 2021; Yamada et al., 2005). We found that heat stress does not drive the release of Atf1 from heterochromatin (Figure 3A), instead the binding affinity between Atf1 and heterochromatin protein Swi6^HP1^ is severely compromised (Figure 4A and 8). This is distinct from the cases in *Drosophila*, mouse and swine, where the release of phosphorylated Atf1 homologues dATF-2 or ATF7 from heterochromatin is the major cause of the disrupted heterochromatin at elevated temperatures (Liu et al., 2019; Seong et al., 2011; Sun et al., 2023).

**Figure 8.**
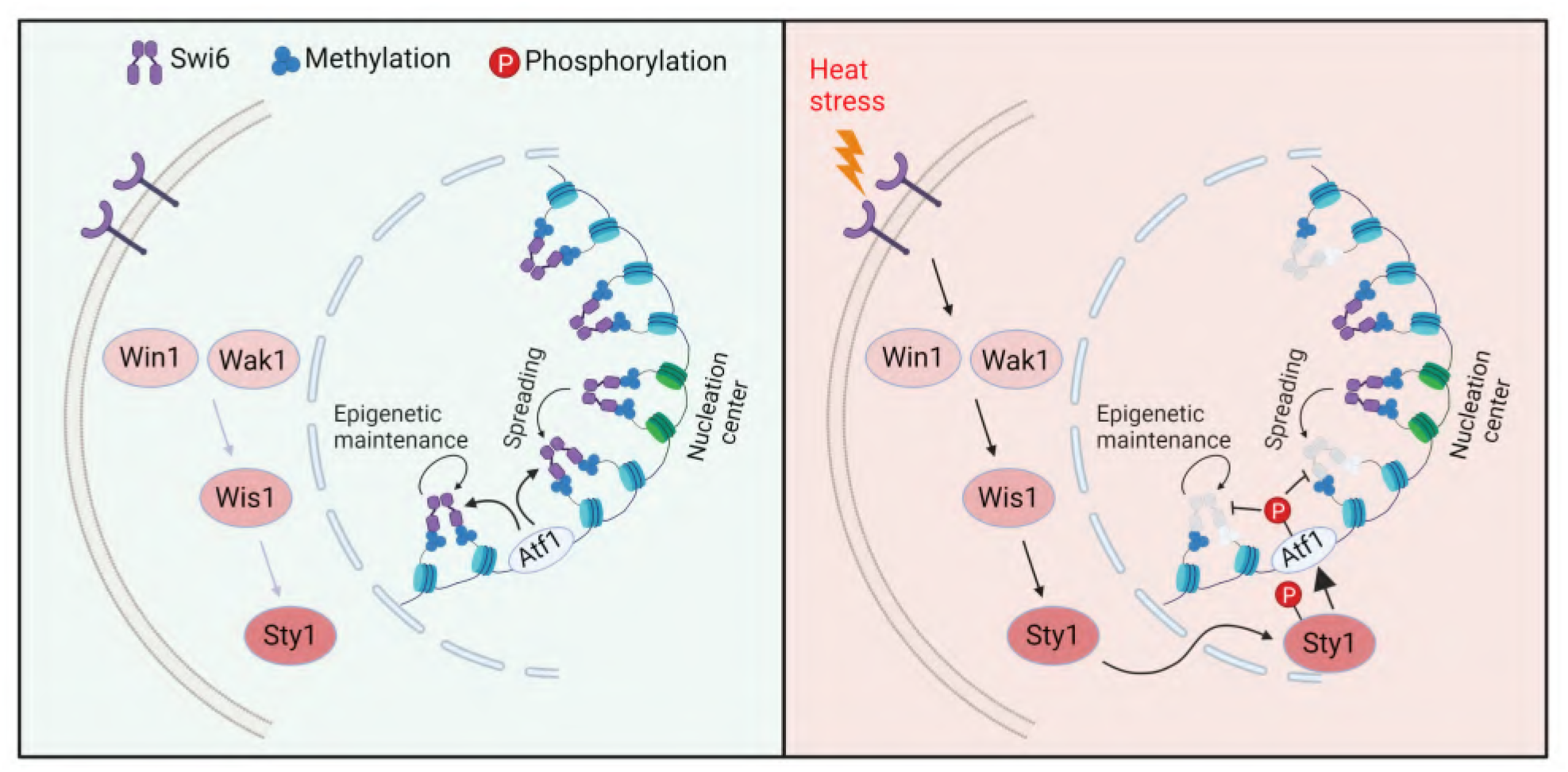
Proposed model for how heat-induced and MAPK-dependent Atf1 phosphorylation provokes epigenetic changes at the *mat* locus in fission yeast. Atf1 plays a dominating role in heterochromatin integrity maintenance and spreading at *mat* locus at normal temperature, but MAPK-mediated Atf1 phosphorylation compromises its binding affinity to Swi6^HP1^, therefore attenuates heterochromatin stability under heat stress.

On the other hand, similarity does exist between fission yeast and higher eukaryotes (such as *Drosophila* and mammals) in that phosphorylation of ATF/CREB family proteins by MAPK are involved in promoting heterochromatin instability. We provided a few lines of evidence to support the idea that phosphorylation of Atf1 by MAPK in *S. pombe* is directly responsible for defective heterochromatin assembly at the *mat* locus under heat stress. First, non-phosphorylatable Atf1 (i.e. Atf1(10A/I)) is indeed more abundantly loaded at both the mating-type region and other Atf1 targets at euchromatic loci than wild type Atf1 (Figure 3F), which efficiently maintains the enrichment of H3K9me3 and Swi6^HP1^ and its binding affinity with Swi6^HP1^ (Figure 3G and 4A). Second, much reduced binding between phosphomimetic Atf1 (i.e. Atf1(10D/E)) and Swi6^HP1^ can be similarly detected between wild type Atf1 and Swi6^HP1^ when MAPK is constitutively activated (i.e. in *wis1-DD* mutants) (Figure 4A and 5E). It still remains mysterious how MAPK-mediated phosphorylation of Atf1 loses its affinity towards Swi6^HP1^, which requires more detailed study in the future.

In previous studies performed in higher eukaryotes, heterochromatin stability has been mostly characterized based on the enrichment of ATF/CREB family protein itself, heterochromatin protein HP1 and H3K9me within examined heterochromatin sites/regions under heat stress. In our current study, in addition to altered binding between Atf1 and Swi6^HP1^ under heat stress, we also found that Clr3, one of the histone modifying enzymes, fails to be efficiently recruited to the *cenH*-flanking sites within the fission yeast *mat* locus (Figure S4B), and *epe1Δ* is able to alleviate silencing defects at this locus (Figure 7A-C). Although we did not observe the weakened binding between Atf1 and Clr3 by *in vitro* pull-down assay (Figure S4A), or enhanced binding of Epe1 at *cenH*-flanking sites (Figure 7D), it does not exclude the possibility that Clr3-recruiting activity of Atf1 is only compromised or Epe1 is more retained respectively in a sub-population of heterochromatic nucleosomes, which cannot be detected by our current methods. It is fairly possible that loss of heterochromatin stability at high temperature may also involves histone modifying enzymes to curb heterochromatin formation in higher eukaryotes.

### Physiological relevance of the loss of heterochromatin stability at *mat* locus in fission yeast

In mammals, the loss of the ATF/CREB family protein-dependent heterochromatin maintenance may cause detrimental consequences. It has been shown that p38-dependent phosphorylation of ATF7 in mice causes its release from the promoters of genes encoding either TERRA (telomere repeat containing RNA) in the sub-telomeric region or Cdk inhibitor p16^Ink4a^, which disrupts heterochromatin and induces TERRA or cellular senescence and a shorter lifespan, respectively (Liu et al., 2019; Maekawa et al., 2019). TERRA can be transgenerationally transmitted to zygotes via sperm and causes telomere shortening in the offspring (Liu et al., 2019). During early porcine embryonic development, high temperatures also trigger the increased expression level of p38, which leads to ATF7 phosphorylation, heterochromatin disruption and eventually the failure of blastocyst formation (Sun et al., 2023).

In fission yeast, the tightly silent mating-type region contributes to the mating-type switching to ensure the presence of almost equal number of opposite mating types cells (Klar, 2007). When exposed to poor nitrogen source conditions, opposite mating type cells mate to form diploid zygotes, and undergo meiosis to form a zygote and ascus containing four spores subsequently (Ohtsuka et al., 2022). Previous studies have shown that the integrity of heterochromatin at the *mat* locus is crucial for efficient mating-type switching in fission yeast (Hansen et al., 2011; Jia et al., 2004b). Indeed, fission yeast cells spend much effort to establish and maintain stable heterochromatin at the *mat* locus by employing multiple mechanisms and factors (Hansen et al., 2011; Jia et al., 2004a; Jia et al., 2004b; Kim et al., 2004; Thon et al., 1999; Wang et al., 2021; Yamada et al., 2005). One very recent study has proposed that the transcription factor Atf1 not only involves in heterochromatin maintenance at the *mat* locus, but also acts in parallel with RNAi machinery and multiple histone-modifying enzymes during heterochromatin establishment steps (Nickels et al., 2022). Even so, fission yeast still suffers the stress-induced defective heterochromatic maintenance at this important locus in its genome. The presence of Atf1 is unable to efficiently fend off phenotypic variation, while its functional disruption can cause even much severe silencing defects at 37°C, as demonstrated by cells with *s1* and *REIII* elements (i.e. the major Atf1 binding sites) deleted (Nickels et al., 2022).

It is generally believed that yeast spores have higher stress tolerance than vegetative cells, which should be helpful for them to survive the unfavorable environment before they meet more friendly conditions. However, it is quite anti-intuitional that fission yeast cells do not mate and therefore fail to undergo meiosis and sporulation at temperatures above 33°C (Brown et al., 2020). The heat stress-induced heterochromatic disruption might interfere with the mating-type switching and reduce mating efficiency, but it seems to be innocuous to vegetatively growing cells. For fission yeast, this feature may be regarded as an “intrinsic flaw” which prevents its use of a better strategy to “escape” from stress and has not been fixed during its evolution. How much heterochromatic maintenance defects at the *mat* locus induced by heat stress is contributing to this feature will be an interesting question for future studies to solve.

## MATERIALS AND METHODS

### Fission yeast strains, media and genetic methods

*Schizosaccharomyces pombe* strains used and created in this study are listed in Table S1. Liquid cultures or solid agar plates consisting of rich medium (YE5S) or minimal medium (PMG5S) containing 4 g/L sodium glutamate as a nitrogen source with appropriate supplements were used as described previously (Forsburg and Rhind, 2006; Moreno et al., 1991). G418 disulfate (Sigma-Aldrich; A1720), hygromycin B (Sangon Biotech; A600230) or nourseothiricin (clonNAT; Werner BioAgents; CAS#96736-11-7) was used at a final concentration of 100 μg/mL where appropriate. Genetic crosses and general yeast techniques were performed as described previously (Forsburg and Rhind, 2006; Moreno et al., 1991).

### Plasmid and yeast strain construction

Yeast strains with C-terminal tagged or deletion of genes were generated by a PCR-based module method with the DNA sequence information obtained from PomBase (https://www.pombase.org). To construct the strain containing *4xtetO-ade6^+^* reporter at the *mat* locus, the vector pBW5/6-4XTetO-ade6^+^ (a kind gift from Robin C. Allshire) was digested with *Pst*I and inserted into the *ura4^+^* locus in strains with *mat3M(Eco*RV*)::ura4^+^*. To construct the plasmid pHBKA81-TetR^0ff^-2xFlag-swi6^ΔCD^, TetR^0ff^-2xFlag was amplified from a vector pDUAL-TetR^0ff^-2xFlag-Stc1 (a kind gift from Robin C. Allshire) and cloned into upstream of *swi6^+^* in the plasmid pHBKA81-swi6-hyg^R^ to generate pHBKA81-TetR^0ff^-2xFlag-swi6. Chromodomain (CD) (80-133aa) of Swi6 was then deleted by Quikgene method (Mao et al., 2011). Finally, the resultant plasmid pHBKA81-TetR^0ff^-2xFlag-swi6^ΔCD^ was linearized by *Apa*I and integrated into the *lys1^+^* locus, generating the strain *lys1Δ::P_adh81_-TetR^off^-2xFlag::hyg^R^*.

### Reporter gene silencing assay

The *ura4^+^* silencing was assessed by growth on PMG5S without uracil or with 1 mg/mL 5-FOA, which is toxic to cells expressing *ura4^+^*. *ade6^+^*silencing was assessed by growth on YE5S with 75 mg/L or 0.5 mg/L adenine, the latter was referred to as YE5S with low adenine. For serial-dilution assays, three serial 10-fold dilutions were made, and 5 μL of each was spotted on plates with the starting cell number of 10^4^.

For *ade6^+^*gene silencing recovery assays, about 500 cells of each strain were plated on YE5S with low adenine medium and incubated at 30℃ or 37℃, the variegated colonies were counted manually. Three variegated colonies grown on solid YE5S with low adenine at 37°C were picked and re-plated on YE5S with low adenine medium and then incubated at 30°C or 37°C to assess the silencing recovery rate.

### Protein extraction and immunoblotting

For total protein extraction, twenty OD_600_ units of *S. pombe* cells at mid-log phase were collected, followed by lysing with glass bead disruption using Bioprep-24 homogenizer (ALLSHENG Instruments, Hangzhou, China) in 200μL lysis buffer containing urea (0.12 M Tris-HCl, pH 6.8, 20% glycerol, 4% SDS, 8 M urea, 0.6 M β-mercaptoethanol).

Western blotting was performed essentially as previously described (Wang et al., 2012). The primary antibodies used for immunoblot analysis of cell lysates were mouse monoclonal anti-Atf1 (abcam, ab18123) (1:2000) and rabbit monoclonal phospho-p38 MAPK (Cell Signaling, 4511T) (1:1000). Cdc2 was detected using rabbit polyclonal anti-PSTAIRE (Santa Cruz Biotechnology, sc-53) (1:10000 dilution) as loading controls. Secondary antibodies used were goat anti-mouse or goat anti-rabbit polyclonal IgG (H+L) HRP conjugates (ThermoFisher Scientific; #31430 or #32460) (1:5000 - 10,000).

### *In vitro* pull-down assay

All recombinant bacterially produced His-Swi6, GST-Clr3, GST-Clr4 and MBP-Clr6 were expressed in *Escherichia coli* BL21 (DE3) cells and purified on Ni^+^ Sepharose^TM^ 6 Fast Flow (for 6His fusions; GE Healthcare, 17531806), Glutathione Sepharose^TM^ 4B (for GST fusions; GE Healthcare, 17075601) or amylose resin (for MBP fusions; New England BioLabs, E8021) according to the manufacturer’s instructions. Yeast cells were lysed by glass bead disruption using Bioprep-24 homogenizer (ALLSHENG Instruments) in NP40 lysis buffer (6 mM Na_2_HPO_4_, 4 mM NaH_2_PO_4_, 1% NP-40, 150 mM NaCl, 2 mM EDTA, 50 mM NaF, 0.1 mM Na_3_VO_4_) plus protease inhibitors. Each purified recombinant protein (about 1 μg) immobilized on resins/beads was incubated with cleared yeast cell lysates for 1 - 3 hours at 4°C. Resins/beads were thoroughly washed and suspended in SDS sample buffer, and then subject to SDS-PAGE electrophoresis and Coomassie brilliant blue (CBB) staining. Western blotting was performed to detect the association between Swi6, Clr3, Clr4 and Clr6 and yeast-expressed Atf1.

### RT-qPCR analysis

Total RNA was extracted using the TriPure Isolation Reagent (Roche). Reverse transcription and quantitative real-time PCR were performed with PrimeScript^TM^ RT Master Mix (Takara, RR037A) and TB Green Premix Ex Taq^TM^ II (Takara, RR820A) in a StepOne real-time PCR system (Applied Biosystems). The relative mRNA level of the target genes in each sample was normalized to *act1^+^*.

### ChIP-qPCR analysis

The standard procedures of chromatin immunoprecipitation were used as previously described (Cam and Whitehall, 2016) with slight modifications. In brief, cells grown in YE5S at 30℃ or 37℃ to mid-log phase were crosslinked with 3% paraformaldehyde for 30 min at room temperature (at 18℃ for Swi6). Thirty OD_600_ units of *S. pombe* cells were collected and lysed by beads disruption using Bioprep-24 homogenizer (ALLSHENG Instruments) in 1 mL lysis buffer (50mM Hepes-KOH pH 7.5, 140 mM NaCl, 1 mM EDTA, 1% Triton X-100, 0.1% deoxycholate) with protease inhibitors. Lysates were sonicated to generate DNA fragments with sizes of 0.5 - 1 kb and immunoprecipitated with mouse monoclonal anti-H3K9me2 (abcam, ab1220), rabbit polyclonal anti-H3K9me3 (abcam, ab8898), mouse monoclonal anti-Atf1 (abcam, ab18123), rabbit polyclonal anti-Swi6 (abcam, ab188276), goat polyclonal anti-myc (abcam, ab9132) or mouse monoclonal anti-Flag M2 (Sigma, F1804), and subjected to pull-down with rProteinA Sepharose^TM^ Fast Flow (GE Healthcare, 17127901) or ProteinG Sepharose^TM^ 4 Fast Flow (GE Healthcare, 17061801). The crosslinking was reversed, and DNA was purified with the ChIP DNA Clean and Concentrator (ZYMO RESEARCH, D5201) kit. Quantitative real-time PCR was performed with TB Green Premix Ex Taq^TM^ II (Takara, RR820A) in a StepOne real-time PCR system (Applied Biosystems). For normalization, serial dilutions of DNA were used as templates to generate a standard curve amplification for each pair of primers, and the relative concentration of target sequence was calculated accordingly. The enrichment of a target sequence in immunoprecipitated DNA over whole-cell extract was calculated and normalized to that of a reference fragment *tub1^+^* as previously described (Wang et al., 2013).

### Statistical analysis

For quantitative analyses of RT-qPCRs and ChIP-qPCR, experiments were repeated three times, and the mean value and standard deviation (s.d.) for each sample was calculated. In order to determine statistical significance of each pair of data, two-tailed unpaired *t*-tests were performed and *p*-values were calculated using GraphPad Prism 7. *p*<0.05 was considered statistically significant.

## Data availability statement

All data are fully available without restriction. All relevant data are within the manuscript figures, supplemental figures, supplemental tables and Source data files.

## ACKNOWLEDGEMENTS

We thank Robin C. Allshire, Marc Buhler, Amikam Cohen, Elena Hidalgo, Genevieve Thon, Janni Petersen and National BioResource Project (NBRP), Japan (http://yeast.nig.ac.jp/yeast/) for fission yeast strains or plasmids; Da-jie Deng for construction of some relevant yeast strains.

## Author contributions

L.S.: investigation, conceptualization, formal analysis, data curation, visualization, writing – original draft preparation; Y.W.: conceptualization, investigation, formal analysis, supervision, writing – review & editing; Q-W.J.: conceptualization, investigation, formal analysis, project administration, supervision, resources, writing – review & editing, funding acquisition.

## Supplemental Figure legends

**Figure S1.**
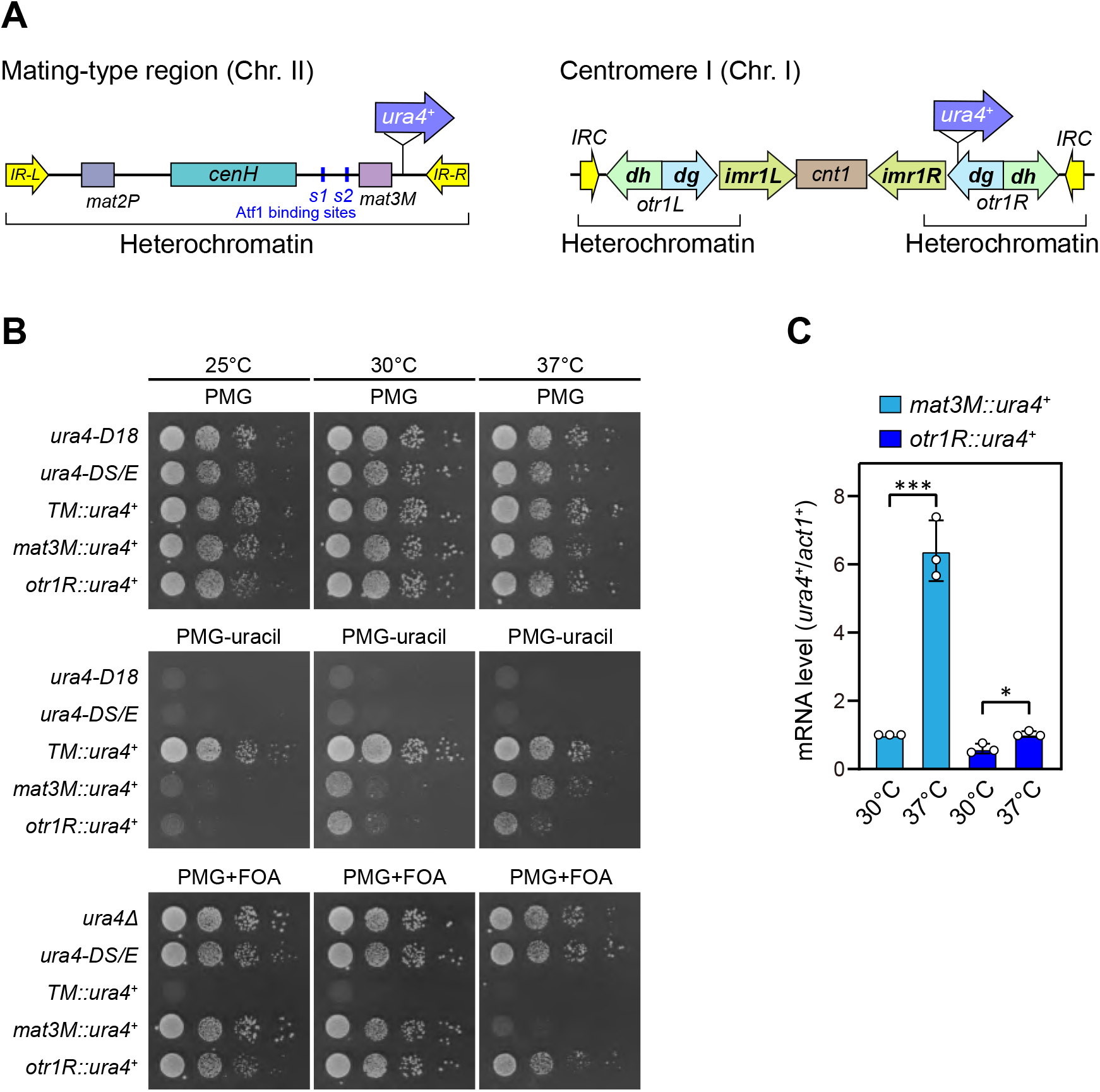
Heat stress leads to defective silencing of reporter *ura4*^+^ at the mating-type region. Related to Figure 1. (A) Schematic of an *ura4^+^* reporter gene inserted into the mating-type region and pericentromeric region of chromosome 2. *cenH*, a DNA element homologous to pericentromeric repeats; *mat2-P* and *mat3-M*, two silent cassettes used for mating-type switching; *IR-L* and *IR-R*, inverted repeats and boundary elements; *s1* and *s2*, two Atf1 binding sites; *cnt1*, central core; *imr1*, innermost repeats; *otr1*, outermost repeats; *dg* and *dh*, tandem repeats in *otr*; *IRC*, inverted repeats and boundary elements. (B) Expression of the *ura4^+^* reporter was monitored by serial dilution spot assay at indicated temperatures. The media used were nonselective PMG5S and selective PMG without uracil or containing 0.15% FOA. *ura4-D18*, complete deletion version of *ura4^+^* gene; *ura4-DS/E*, truncated version of *ura4^+^* gene; *TM::ura4^+^*, *ura4^+^* gene inserted at a random site within euchromatin region in the genome. (C) RT-qPCR analyses of *ura4^+^* reporter. The relative *ura4^+^* mRNA level was quantified with a ratio between *mat3M::ura4^+^* and *act1^+^* in 30°C samples being set as 1.00. Error bars indicate mean ± standard deviation of three independent experiments. Two-tailed unpaired *t*-test was used to derive *p* values. \**p*<0.05; \*\*\**p*<0.001.

**Figure S2.**
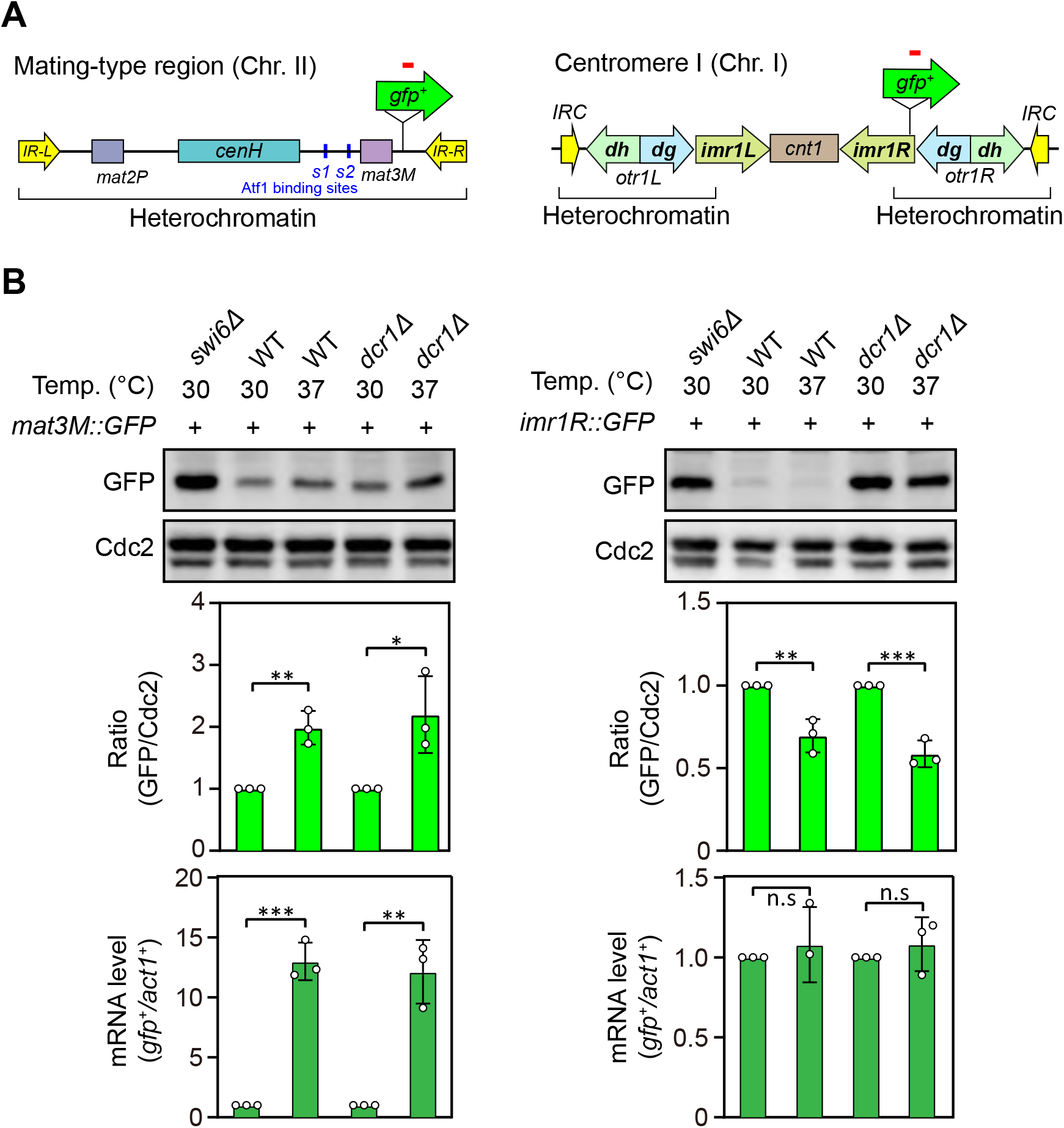
Heat stress leads to defective silencing of *gfp*^+^ reporter gene at the mating-type region. Related to Figure 1. (A) Schematic of a *gfp^+^* reporter gene inserted into the mating-type region (*mat3M*) and pericentromeric region (*imr1R*). (B) Western blotting and RT-qPCR analyses were used to measure expression of the *gfp^+^* reporter. (*Upper*) Western blotting analyses of the protein level of GFP in wild type, *dcr1Δ* and *swi6Δ* cells at 30°C and 37°C. (*Middle*) Quantitative analyses of the protein level of GFP in wild type and *dcr1Δ* cells at 30°C and 37°C. (*Lower*) RT-qPCR analyses of *gfp^+^* reporter in wild type and *dcr1Δ* cells at 30°C and 37°C. For quantifications, the relative protein or mRNA levels were quantified with a ratio between GFP and Cdc2 or between transcripts of *gfp^+^* and *act1^+^* in 30°C samples being set as 1.00, respectively. Error bars indicate mean ± standard deviation of three independent experiments. Two-tailed unpaired *t*-test was used to derive *p* values. n.s, not significant; \**p*<0.05; \*\**p*<0.01; \*\*\**p*<0.001.

**Figure S3.**
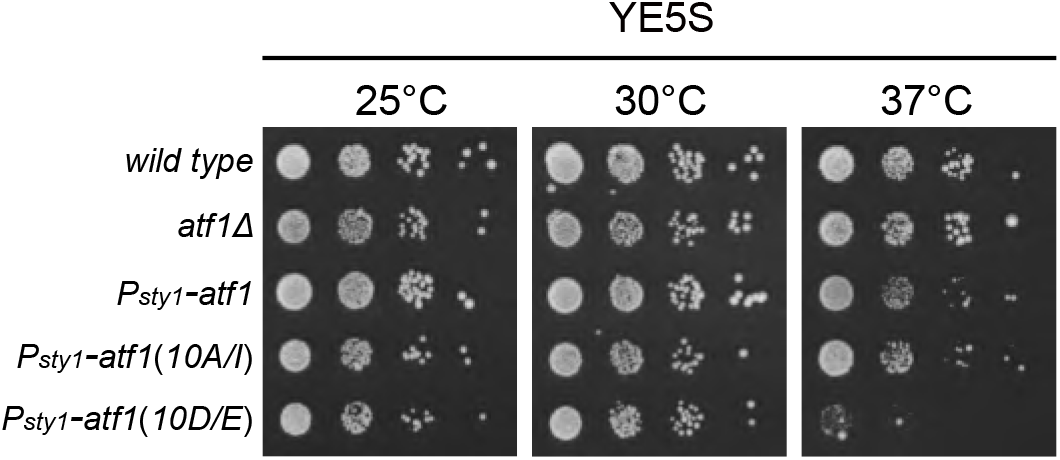
Expression of Atf1(10D/E) leads to lethality at 37°C. Yeast strains with indicated genotypes were first grown in liquid YES at 25°C, then spotted onto YES plates. Plates were incubated at indicated temperatures for > 3 days.

**Figure S4.**
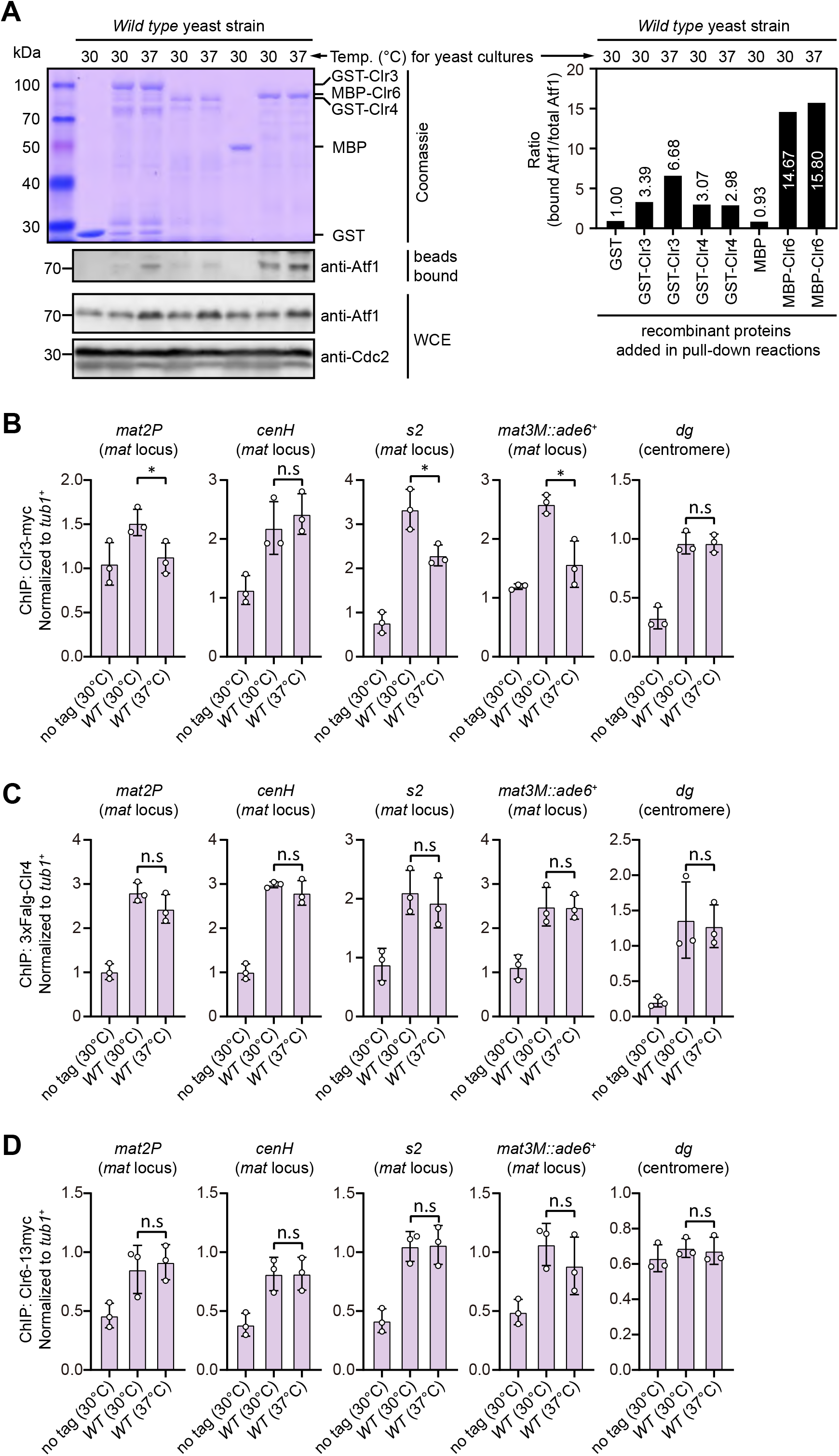
The association between Clr3, Clr4 or Clr6 and Atf1 analyzed by *in vitro* binding assay, and their enrichment at different heterochromatic regions under heat stress analyzed by ChIP-qPCR. (A) *In vitro* binding assays for binding affinity between Atf1 and Atf1-associated heterochromatin factors Clr3, Clr4 or Clr6. Cell lysates prepared from wild type yeast strain grown at either 30°C or 37°C were incubated with bacteria-expressed GST-Clr3, GST-Clr4 or MBP-Clr6. (*Left*) Bound and total Atf1 were detected by immunoblotting with Cdc2 used as a loading control. Results are representative of three independent experiments. (*Right*) The relative binding affinity between Atf1 and Clr3, Clr4 or Clr6 was quantified with a ratio between GST and Atf1 from a 30°C culture being set as 1.00. (B-D) ChIP-qPCR analyses of Clr3 (B), Clr4 (C) or Clr6 (D) levels at representative heterochromatic loci in wild type cells grown at 30°C and 37°C. Relative enrichment of Clr3, Clr4 or Clr6 was normalized to that of a *tub1^+^* fragment. Error bars represent standard deviation of three experiments. Two-tailed unpaired *t*-test was used to derive *p* values. n.s, not significant; \**p*<0.05; \*\**p*<0.01; \*\*\**p*<0.001.

---

**Table S1.**
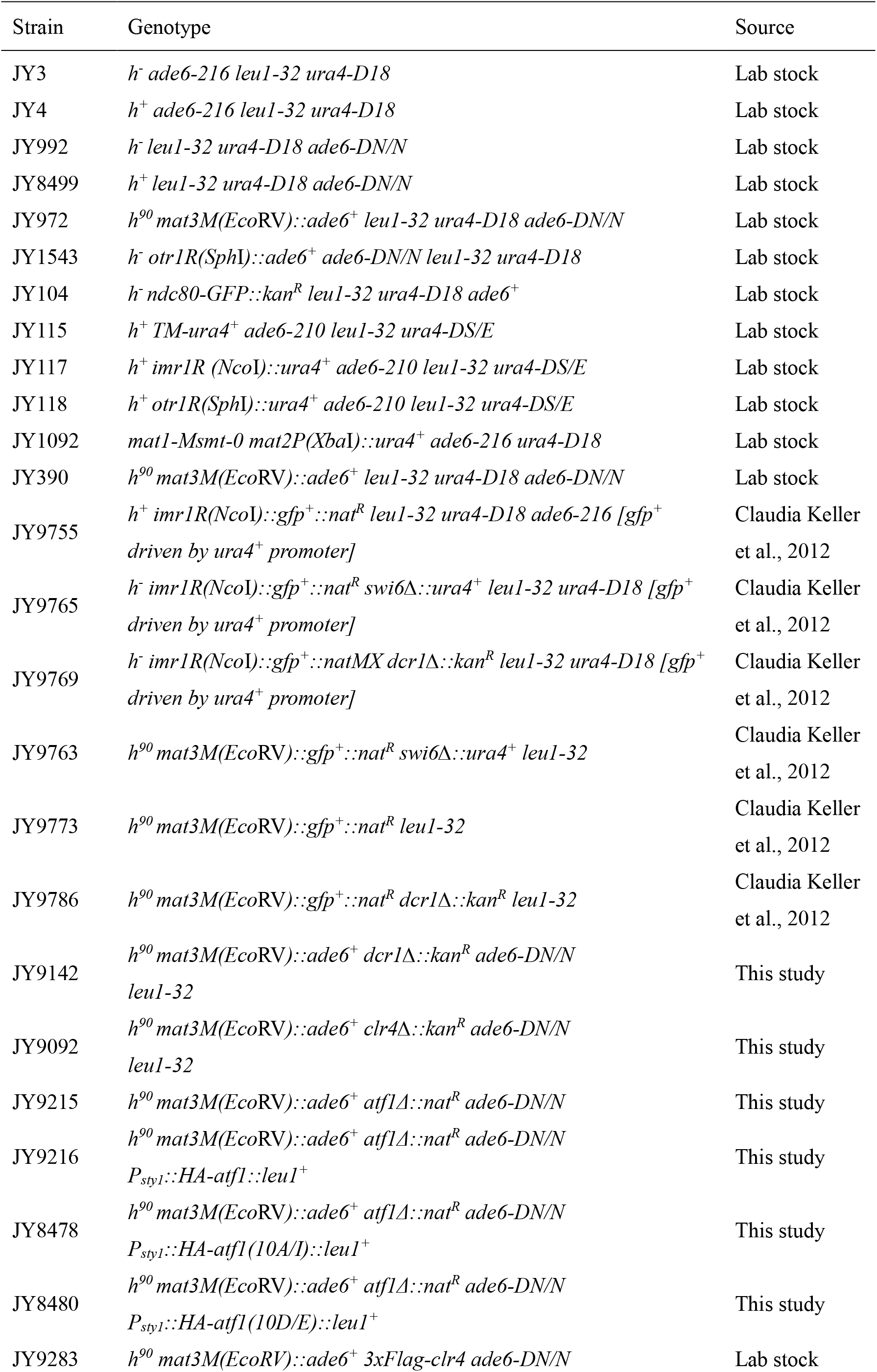
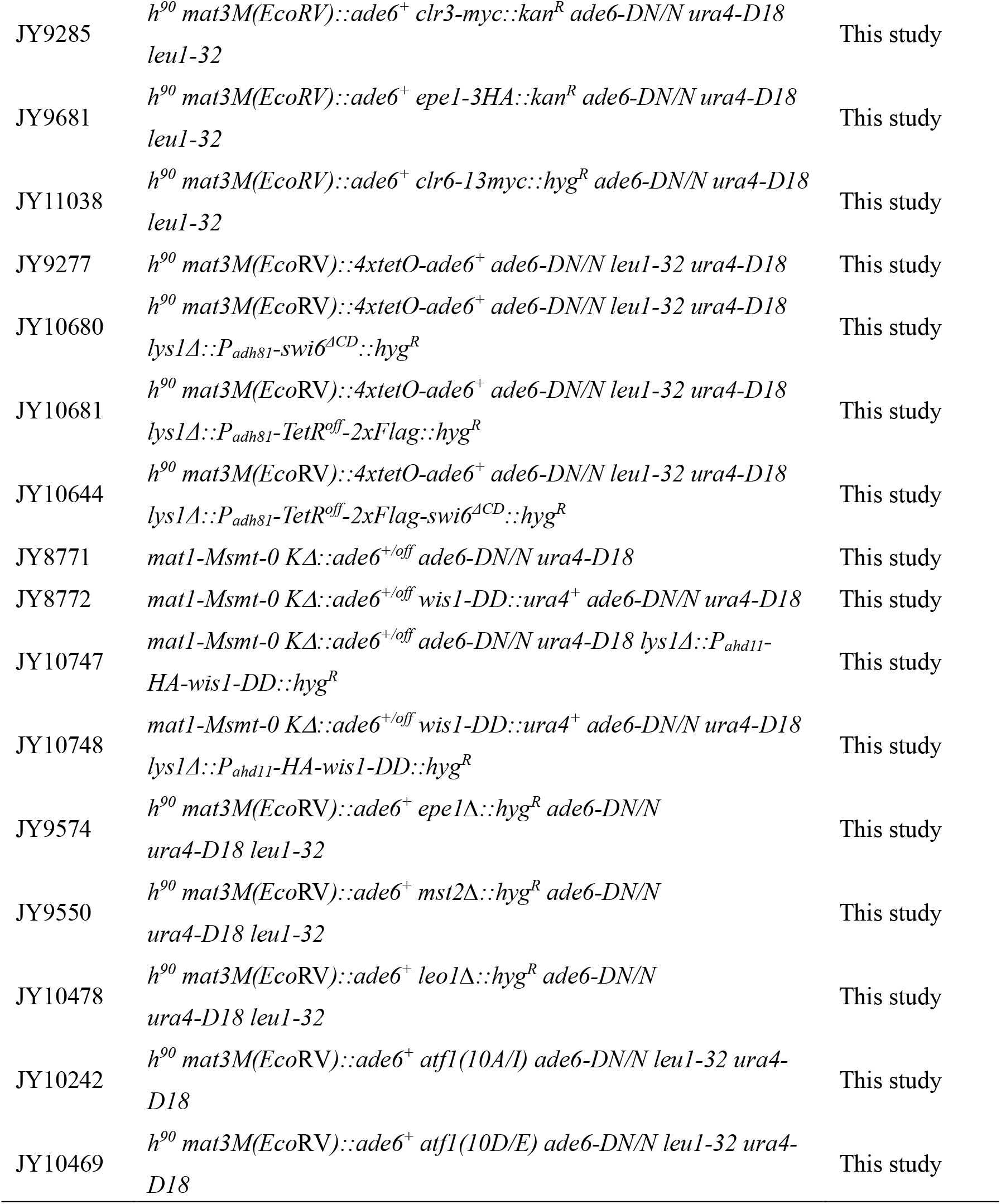
Yeast strains used in this study.

**Table S2.**
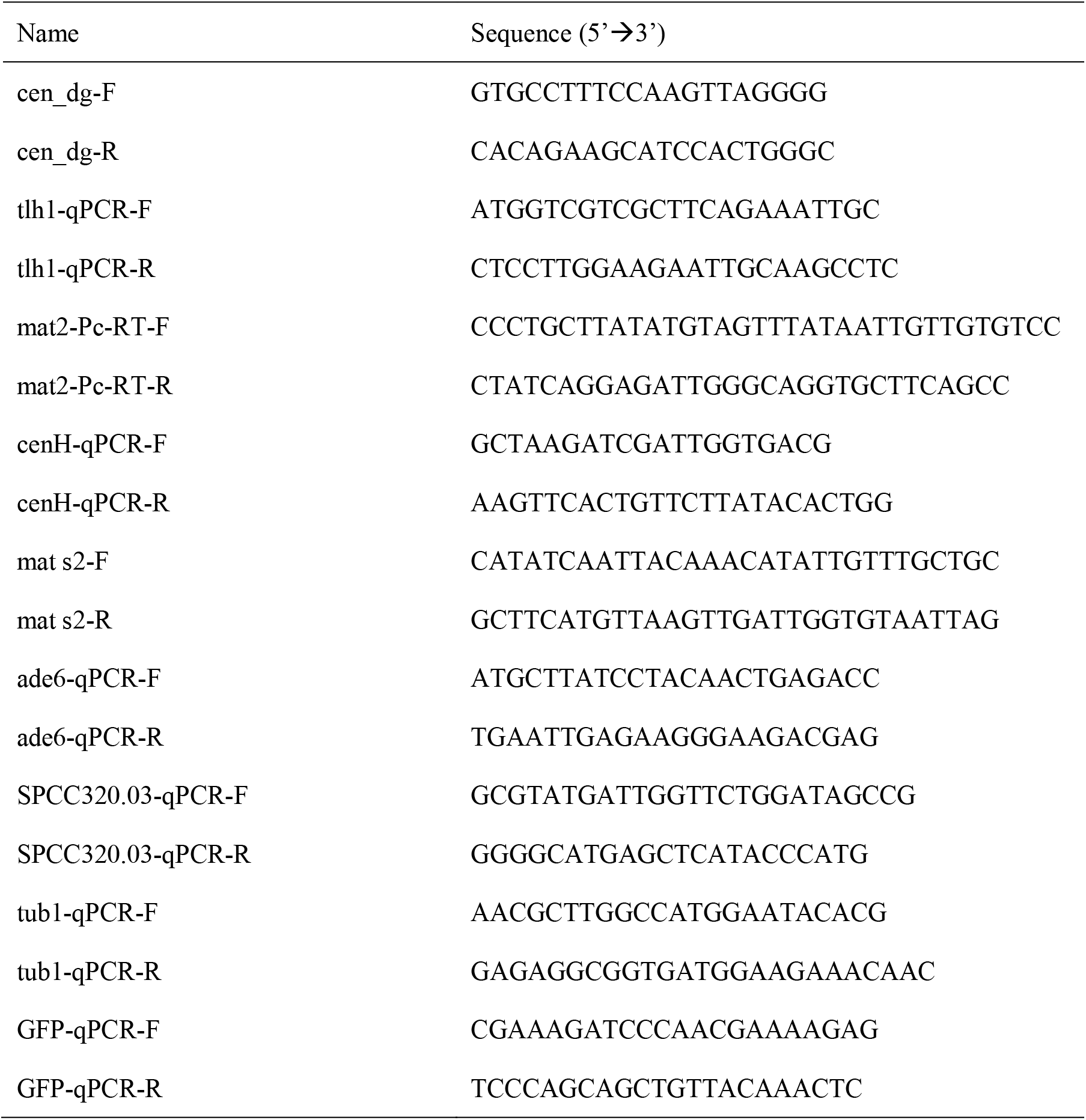
Primers used for RT-qPCR and qPCR.

## Source Data Legends

Figure 1-Source Data [raw data of colony color assay, RT-qPCR, H3K9me2/3 ChIP]

Figure 2-Source Data [raw data of colony color assay, RT-qPCR, H3K9me3 ChIP]

Figure 3-Source Data 1 [raw data of colony color assay, RT-qPCR, Atf1/Swi6/H3K9me3 ChIP]

Figure 3-Source Data 2 [full raw unedited blot (Atf1) of Figure3C]

Figure 3-Source Data 3 [full raw unedited blot (Cdc2) of Figure3C]

Figure 3-Source Data 4 [uncropped blots of Figure 3C]

Figure 4-Source data 1 [raw data of Swi6 ChIP]

Figure 4-Source Data 2 [full raw unedited Coomassie gel (His-Swi6) of Figure 4A]

Figure 4-Source Data 3 [full raw unedited blot (bead bound-Atf1) of Figure 4A]

Figure 4-Source Data 4 [full raw unedited blot (WCE-Atf1) of Figure 4A]

Figure 4-Source Data 5 [full raw unedited blot (WCE-Cdc2) of Figure 4A]

Figure 4-Source Data 6 [uncropped blots of Figure 4A]

Figure 5-Source Data 1 [raw data of RT-qPCR, Swi6/H3K9me3 ChIP]

Figure 5-Source Data 2 [full raw unedited blot (Sty1-P) of Figure 5D]

Figure 5-Source Data 3 [full raw unedited blot (Atf1) of Figure 5D]

Figure 5-Source Data 4 [full raw unedited blot (Cdc2) of Figure 5D]

Figure 5-Source Data 5 [uncropped blots of Figure 5D]

Figure 5-Source Data 6 [full raw unedited gel (Coomassie) of Figure 5E]

Figure 5-Source Data 7 [full raw unedited blot (beads bound-Atf1) of Figure 5E]

Figure 5-Source Data 8 [full raw unedited blot (WCE-Atf1 and Cdc2) of Figure 5E]

Figure 5-Source Data 9 [uncropped gel and blots of Figure 5E]

Figure 6-Source Data [raw data of colony color assay, RT-qPCR, TetR-Flag/H3K9me3 ChIP]

Figure 7-Source Data [raw data of colony color assay, RT-qPCR, H3K9me3/Epe1-3HA ChIP]

Figure S1-Source Data [raw data of RT-qPCR]

Figure S2-Source Data 1 [raw data of GFP level measurement, RT-qPCR]

Figure S2-Source Data 2 [full raw unedited blot (mat3M-GFP) of Figure S2B]

Figure S2-Source Data 3 [full raw unedited blot (Cdc2) of Figure S2B]

Figure S2-Source Data 4 [full raw unedited blot (imr1R-GFP) of Figure S2B]

Figure S2-Source Data 5 [full raw unedited blot (Cdc2) of Figure S2B]

Figure S2-Source Data 6 [uncropped blots of Figure S2B]

Figure S4-Source Data 1 [raw data of *in vitro* binding assay, Clr3/Clr4/Clr6 ChIP]

Figure S4-Source Data 2 [full raw unedited Coomassie gel of Figure S4A]

Figure S4-Source Data 3 [full raw unedited blot (bead bound-Atf1) of Figure S4A]

Figure S4-Source Data 4 [full raw unedited blot (WCE-Atf1) of Figure S4A]

Figure S4-Source Data 5 [full raw unedited blot (WCE-Cdc2) of Figure S4A]

Figure S4-Source Data 6 [uncropped blots of Figure S4A]

